# Antagonistic Neural Circuits Drive Opposing Behaviors towards the Young in Females

**DOI:** 10.1101/2023.03.12.532288

**Authors:** Long Mei, Rongzhen Yan, Luping Yin, Regina Sullivan, Dayu Lin

## Abstract

In many species, including mice, females show strikingly different pup-directed behaviors based on their reproductive state^1,2^. Naïve wild female mice often kill pups while lactating females are dedicated to pup caring^3,4^. The neural mechanisms that mediate infanticide and its switch to maternal behaviors during motherhood remain unclear. Here, based on the hypothesis that maternal and infanticidal behaviors are supported by distinct and competing neural circuits^5,6^, we used the medial preoptic area (MPOA), a key site for maternal behaviors^7–11^, as a starting point and identified three MPOA-connected brain regions that drive differential negative pup-directed behaviors. Further functional manipulation and *in vivo* recording revealed that estrogen receptor alpha (Esr1) expressing cells in the principal nucleus of the bed nucleus of stria terminalis (BNSTpr^Esr1^) are necessary, sufficient, and naturally activated during infanticide in female mice. Furthermore, MPOA^Esr1^ and BNSTpr^Esr1^ neurons form reciprocal inhibition and change their excitability in opposite directions with reproductive state. The shift in balance between BNSTpr^Esr1^ and MPOA^Esr1^ cell activity is likely a key mechanism for the behavioral switch during motherhood.

## Introduction

At birth, for nearly all mammalian species, the young are vulnerable and powerless. Its chance of survival critically depends on care and protections from the parents, especially mothers. Consequently, a set of robust and stereotypical maternal behaviors have evolved to ensure the needs of the young are met. In mice, postpartum females spend the vast majority of their time in the nest to nurse and groom and lick pups to keep them clean and stimulate urination and defecation^9,12^. When pups wander from the nest and call for help, mothers quickly search for and retrieve them back to the nest. However, females do not always care for pups. Across a wide range of mammalian species, it is not uncommon for virgin females to show hostile behaviors towards pups of the same species^13,14^. From an evolutionary perspective, this behavior is believed to occur in males and females to increase the killer’s reproductive success by freeing up critical resources for her future progeny, including food, shelters, care and a social position^13,15,16^. In a survey involving 289 mammalian species, infanticide was found in 31% of species, with a higher percentage in species that breed in groups^13^. Despite the prevalence of infanticide in mammalian females, including mice^3,4^, it is rarely studied under laboratory conditions partly because adult females of many inbred strains of mice, e.g., C57BL/6, rarely show such behavior, likely due to inbreeding^17,18^. Female mice of outbred strains, e.g. Rockland-Swiss, appear to have retained more naturalistic behaviors, including a higher level of infanticide than inbred mice, although the exact likelihood varies with age^19^.

The neural circuit of maternal behaviors has been extensively studied and medial preoptic area (MPOA), an evolutionary conserved region situated at the anterior end of the hypothalamus, has been firmly established as a key region for maternal behaviors through numerous functional and *in vivo* recording experiments since 1970s^7–11^. Lesion studies first revealed that damage to MPOA impairs maternal behaviors, in extreme cases, turning caring to killing^20,21^. Later pharmacological, pharmacogenetic and optogenetic inactivation all corroborate these lesion results^22–26^. Conversely, estrogen supplement or optogenetic activation of the MPOA cells that express estrogen receptor alpha (Esr1) or galanin facilitates various aspects of maternal behaviors, such as pup retrieval and grooming^23–29^. Consistent with the functional results, *in vivo* electrophysiological recording and imaging revealed that subpopulations of MPOA neurons are activated during different components of maternal behaviors^23,25,26^. Interestingly, MPOA cells that are relevant for parental behaviors are mainly inhibitory. 75% of Esr1 and 90% galanin expressing cells are GABAergic^24,26,30^ and a recent study found that activating GABAergic cells in the MPOA is sufficient to elicit pup retrieval and nest building^31^ whereas MPOA glutamatergic cells are activated by aversive stimuli and mediate anxiety-like behaviors^32^.

In contrast to our extensive knowledge in the maternal circuit, very little is known regarding the neural substrates responsible for infanticide in females. A few recent studies started to reveal brain regions relevant for infanticide in males but their roles in females are either minimal or unexplored^33–36^. Yousuke et. al. showed that rhomboid nucleus of the bed nucleus of stria terminalis (BNSTrh) express high level of c-Fos after infanticide in male mice^33^. Lesion in the BNSTrh suppresses infanticide in virgin males^33^. Chen et. al. found that GABAergic cells in medial amygdala posterodorsal part (MeApd) drives infanticide in male but not female mice^34^. Autry et. al. reported that inhibition of perifornical area urocortin-3 cells suppresses infanticide in males while activating the cells induced infanticide in a small percentage of females^36^. Most recently, Sato et. al. showed that pharmacogenetic activation of amygdalohippocampal area (AHi) oxytocin receptor (OXTR) expressing cells that project to MPOA promotes infant-directed attack in fathers but whether the same manipulation will cause infanticide in mothers was not tested^35^.

Thus, the neural circuit that drives hostile pup-directed behaviors in females is virtually unknown. Given that MPOA cells are largely inhibitory and damage to MPOA can lead to infanticide, we hypothesize that maternal care circuit and infanticide circuit are reciprocally connected and counteract one another. Based on this hypothesis, we systematically manipulated regions that are directly connected with MPOA and identified multiple brain areas that robustly promote negative pup-directed behaviors in female mice. We then further examined one of the regions, the principal nucleus of the bed nucleus of stria terminalis (BNSTpr), and revealed its indispensable role in infanticide, its reciprocal inhibitory connection with MPOA^Esr1^ cells, and its maternal state dependent changes in physiological properties.

## Results

### Identification of brain regions that promote negative pup-directed behaviors using MPOA as an entry point

Consistent with the previous reports, we found that infanticide is rare in adult C57BL/6 females^17,18,37^. It was only observed in 2 out of 165 virgin C57BL/6 female mice tested in our study. In contrast, approximately one third (50/146) of virgin Swiss Webster (SW) females attacked and killed pups, making SW mice a suitable animal model for studying the neural mechanisms underlying naturally occurring infanticide in the lab.

Based on the hypothesis that infanticide and maternal circuits form mutual inhibition, we predict that regions relevant for infanticide are likely connected to MPOA, a central region for maternal care. To identify these regions, we injected 50 nl high titer (>1 × 10^13^) AAV1 expressing hSyn-Cre into MPOA of Ai6 SW female mice^38^ (**Extended Data Fig. 1a**). High titer AAV1 travels both anterogradely^39^ and retrogradely^40^ to label downstream and upstream cells. During the surgery, 50 nl AAV2 expressing Cre-dependent mCherry was also co-injected to label the primary injection site. Three weeks after virus injection, GFP^+^ cells were found to be widely distributed, including 18 regions that each contained more than 1% of total labeled cells (**Extended Data Figs. 1b and 1c**). Half (9/18) of the highly connected regions were within the hypothalamus and collectively contributed to 69% of total labeled cells (**Extended Data Fig. 1c**). Outside of the hypothalamus, the densely labeled cells were found in lateral septum (LS) and nucleus accumbens (NAc), paraventricular thalamus (PVT), BNSTpr, medial amygdala posterodorsal part (MeApd), posterior amygdala (PA), ventral subiculum (SUBv), ventral tegmental area (VTA) and periaqueductal gray (PAG) (**Extended Data Fig. 1c**). Among those areas, BNSTpr, PVT and MeApd showed a significantly higher number of Fos^+^ cells in hostile females after infanticide in comparison to non-hostile females after pup interaction (**Figs. 1a-1b and Extended Data Figs. 1d-1g**). When considering only Fos^+^ cells that are connected with MPOA, we found that BNSTpr, MeApd and supramammillary nucleus (SUM) had significantly more double positive cells in hostile females than control females, while paraventricular nucleus (PVN), LS and ventrolateral part of the ventromedial hypothalamus (VMHvl) showed a trend towards increase (**Fig. 1c and Extended Data Figs. 1d-1g**).

**Fig. 1:**
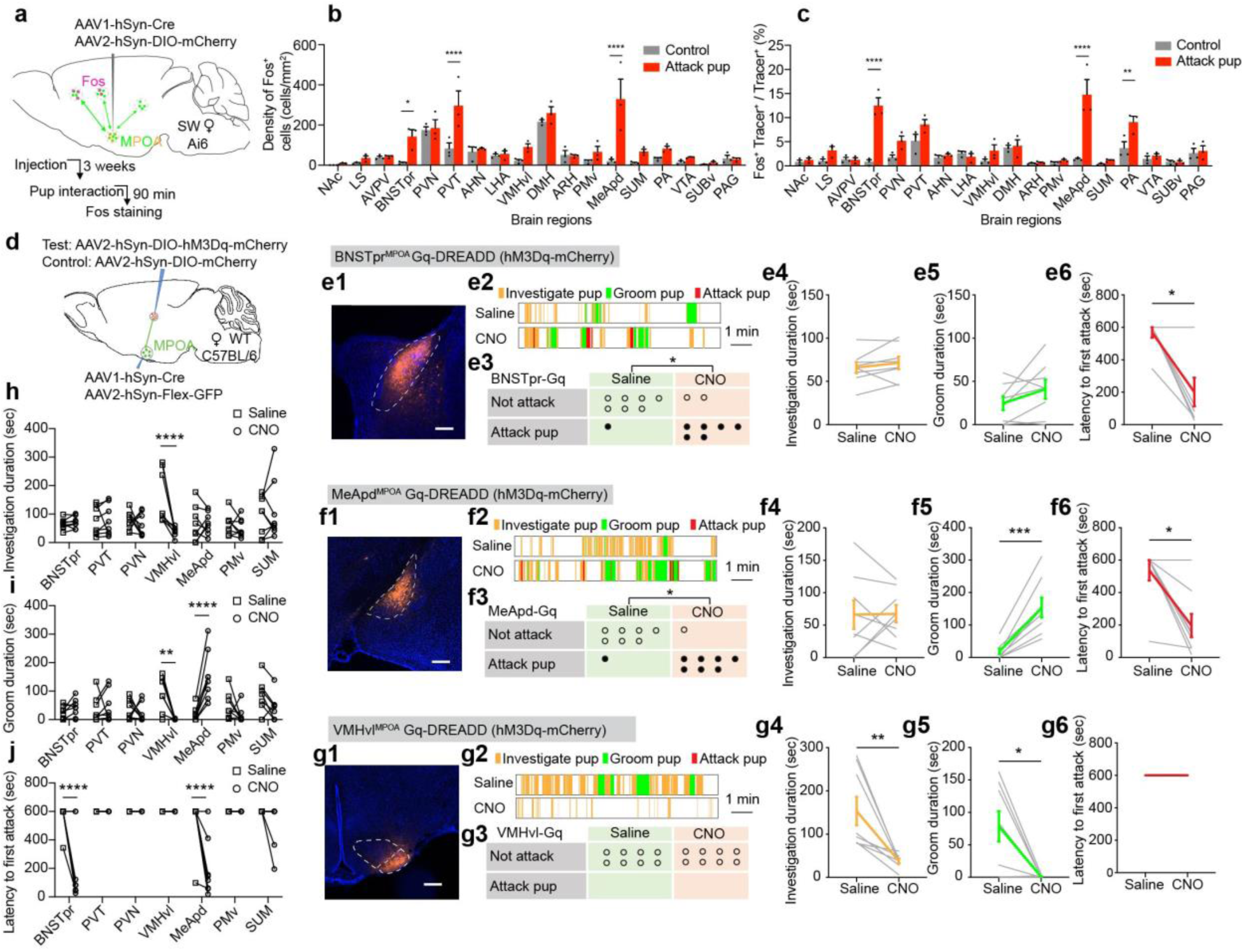
Functional screening of brain regions relevant for negative pup-directed behaviors in females. **(a)** Experimental design to identify MPOA-connected cells that are activated by infanticide in female mice. **(b)** The density of c-Fos expressing cells in various MPOA-connected brain regions in control and infanticidal female mice. n = 3 mice for each group. Two-way ANOVA with Bonferroni’s multiple comparisons test. *p < 0.05, ****p < 0.0001, Error bars: SEM. **(c)** The percentage of MPOA-connected cells that express c-Fos in infanticide and control females. n = 3 mice for each group. Two-way ANOVA with Bonferroni’s multiple comparisons test. **p < 0.01, ****p < 0.0001. Error bars: SEM. **(d)** Experimental design to chemogenetically activate MPOA-connected regions. **(e1, f1, g1)** Representative histology images showing virally expressed hM3Dq-mCherry (red) in BNSTpr^MPOA^ (**e1**), MeApd^MPOA^ (**f1**), VMHvl^MPOA^ (**g1**) cells. Scale bars: 200 µm. **(e2, f2, g2)** Representative raster plots showing pup-directed behaviors after saline (top) or CNO (bottom) injection in BNSTpr^MPOA^ (**e2**), MeApd^MPOA^ (**f2**), VMHvl^MPOA^ (**g2**) mice. Each behavior test session lasts 10 min. **(e3, f3, g3)** Number of female mice that show attack or not after saline or CNO injection in BNSTpr^MPOA^ (**e3**), MeApd^MPOA^ (**f3**), VMHvl^MPOA^ (**g3**) mice. Each dot represents one animal. Fisher’s exact test. *p < 0.05. **(e4-e6, f4-f6, g4-g6)** Investigating pup duration (**e4, f4, g4**), grooming pup duration (**e5, f5, g5**), and latency to attack pups (**e6, f6, g6**) after saline or CNO injection in BNSTpr^MPOA^ (**e4-e6**), MeApd^MPOA^ (**f4-f6**), VMHvl^MPOA^ (**g4-g6**) mice. Each gray line represents one animal. Colored lines represent mean ± SEM. t-tests, **p < 0.05, **p < 0.01, ***p < 0.001. **(h-j)** Summary of changes in pup investigation duration (**h**), pup grooming duration (**I**) and latency to attack pups (**j**) after CNO injection in comparison to saline injection in animals with hM3Dq-mCherry expression in MPOA-connected cells in various brain regions. Two-way ANOVA with Bonferroni’s multiple comparisons test. **P < 0.01, ****P < 0.0001. n = 8 mice for each group. Source data provided. Details of the statistical analyses and sample sizes can be found in Statistic Summary Table.

Based on the density of Fos^+^ cells induced by infanticide and the cell connectivity with MPOA, we decided to systematically activate MPOA-connecting cells in the BNSTpr, MeApd, SUM, PVN, PVT and VMHvl and examine changes in pup-directed behavior. We also included PMv in our manipulation list given its function in inter-male aggression^41,42^. For each animal, we injected 100 nl high titer AAV1 expressing hSyn-Cre bilaterally into the MPOA and Cre-dependent hM3Dq-mCherry into one of the targeting regions in C57BL/6 wildtype female mice (**Fig. 1d**). During the surgery, AAV2 expressing Cre-dependent GFP was also injected into the MPOA and only animals with GFP expression centered in the MPOA were included for analysis (**Fig. 1d**). As we aimed towards identifying regions that enhance infanticide, we used C57BL/6 female mice for this experiment given their near zero spontaneous infanticide.

Three weeks after virus injection, we injected saline or CNO (1 mg/kg) intraperitoneally into the test mouse and 30 minutes later introduced three 1-4 days old pups into the home cage for 10 minutes. Strikingly, pharmacogenetic activation of MPOA-connecting BNSTpr (BNSTpr^MPOA^) or MeApd (MeApd^MPOA^) cells elicited repeated attack towards pups in vast majority of tested C57BL/6 females (**Figs. 1e, and 1f**) while the total duration of pup investigation did not change (**Figs. 1e4, and 1f4**). Activating the MeApd^MPOA^ cells, but not BNSTpr^MPOA^ cells, also increased the duration of pup grooming (**Figs. 1e5 and 1f5**), a behavioral change that was also observed upon optogenetic activation of MeApd GABAergic cells^34^. Animals that expressed mCherry in the MeApd^MPOA^ or BNSTpr^MPOA^ cells showed no behavioral change after CNO injection and none of the mCherry control animals showed infanticide (**Extended Data Figs. 2a and 2b**).

When the MPOA-connecting VMHvl (VMHvl^MPOA^) cells were pharmacogenetically activated, the test females showed a general avoidance towards the pups. They spent significantly less time investigating or grooming the pups. However, none of the VMHvl^MPOA^ activated females showed infanticide (**Fig. 1g**). When the MPOA-connecting PVT, PVN, PMv and SUM cells were activated, we observed no significant behavior change towards the pups (**Extended Data Figs. 2c-2f**).

These results suggest that pup-directed behaviors, including pup avoidance, pup grooming and infanticide, are mediated by different combinations of brain regions (**Figs. 1h-1j**). BNSTpr^MPOA^ cells may play an important and specific role in driving infanticide during pup encounter.

### Optogenetic activation of BNSTpr^MPOA^ neurons elicits infanticide in virgin females

To understand whether the BNSTpr^MPOA^ cells drive infanticide acutely or only increase the likelihood of its expression, we optogenetically activated BNSTpr^MPOA^ cells bilaterally in virgin C57BL/6 female mice (**Fig. 2a**). During testing, we introduced several pups into the female home cage and delivered a serial of laser pulses (20 ms, 20 Hz, 20 s) with gradually increasing laser intensity (0.5 − 5 mW) (**Figs. 2b and 2c**). Upon light stimulation at even the lowest intensity, 10/11 ChR2-expressing females showed time-locked pup attack in approximately 65% stimulation trials while none of the GFP-expressing females attacked the pups **(Figs. 2d-2g**). Once the animal encountered the pup, the attack was initiated with an average latency of approximately 3 s, suggesting that activation of BSNTpr^MPOA^ cells drive infanticide acutely (**Fig. 2h**). Increasing light intensity did not change the induced behavior qualitatively although the percentage of infanticide-induced trials showed a trend of decrease at high light intensity (> 3 mW) (**Fig. 2g**).

**Fig. 2:**
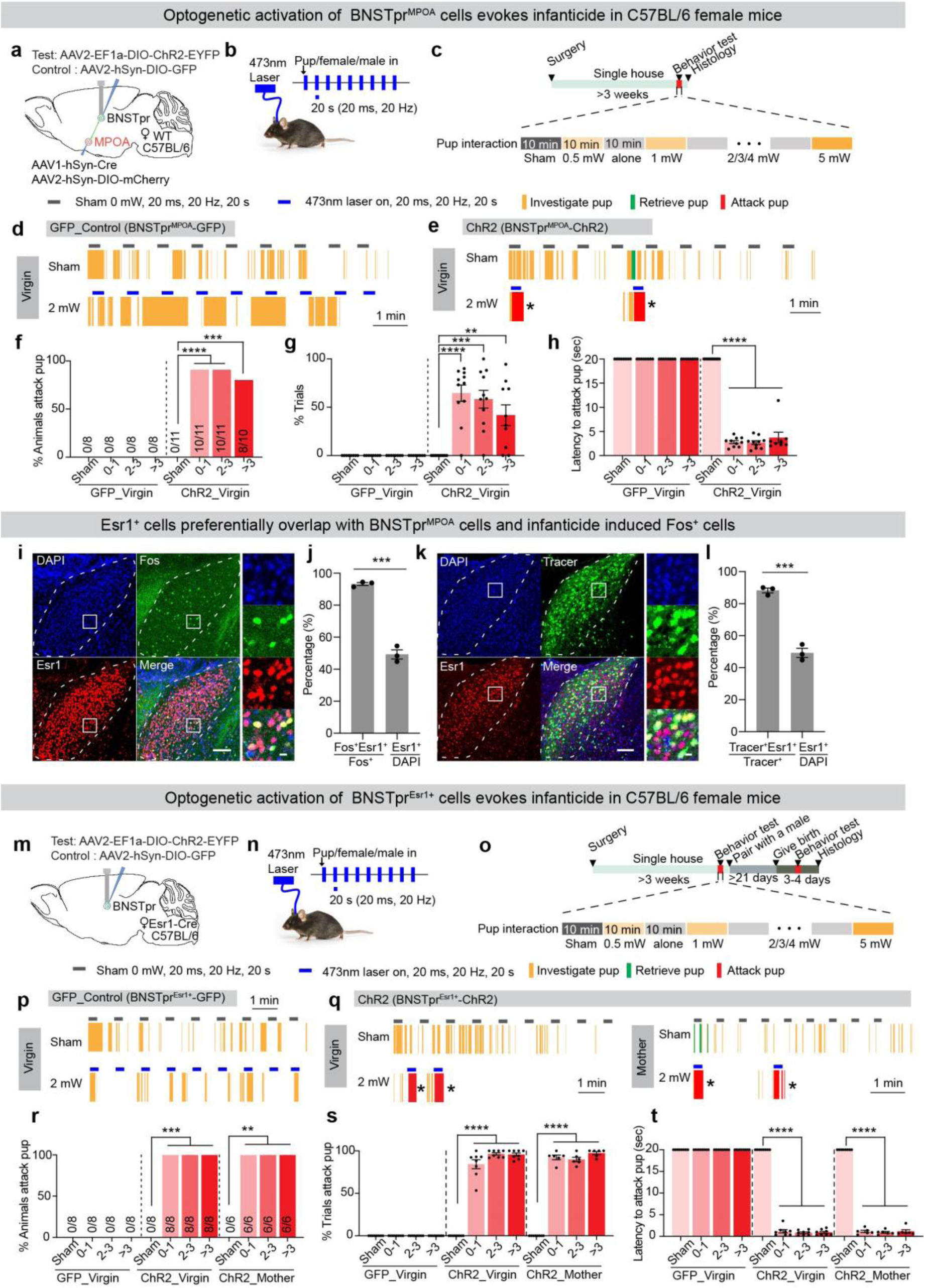
Optogenetic activation of BNSTpr^MPOA^ or BNSTpr^Esr1^ neurons elicits infanticide in females. **(a)** Experimental design to optogenetically activate BNSTpr^MPOA^ cells. **(b)** Light delivery protocol. **(c)** Experimental timeline. **(d and e)** Representative raster plots showing pup-directed behaviors during sham and 2 mW light stimulation in virgin female mice expressing GFP (**d**) or ChR2-eYFP (**e**) in BNSTpr^MPOA^ cells. *Remove hurt pups. **(f)** Percentage of animals that attacked pups during sham or light delivery. Fisher’s exact test. ***P < 0.001, ****P < 0.0001. n = 8 mice for GFP control, n = 11 mice for ChR2 group. **(g)** Percentage of trials showing pup attack. Each dot represents one mouse. Error bars ± SEM. Two-way ANOVA with Bonferroni’s multiple comparisons test. **P < 0.01, ***P < 0.001, ****P < 0.0001. n = 8 mice for GFP control, n = 11 mice for ChR2 group. **(h)** Averaged latency to attack pup upon encountering the pup following sham or light stimulation. Error bars ± SEM. Two-way ANOVA with Bonferroni’s multiple comparisons test. ****P < 0.0001. n = 8 mice for GFP control, n = 11 mice for ChR2 group. **(i)** Representative images showing overlap between Esr1 (red) and infanticide induced c-Fos (green) in the BNSTpr. Right shows the enlarged view of the boxed area. Scale bars: 100 µm (left) and 10 µm (right). **(j)** The percentage of infanticide induced c-Fos positive cells that express Esr1 is significantly higher than the percentage of total BNSTpr cells that express Esr1. n = 3 mice. Error bars ± SEM. Unpaired T-test. ***p < 0.001. **(k)** Images showing the overlap between Esr1 (red) and zsGreen (green) in the BNSTpr in an Ai6 animals injected with AAV1-hSyn-Cre at the MPOA. Right shows the enlarged view of the boxed area. Scale bars: 100 µm (left) and 10 µm (right). **(l)** The percentage of MPOA-connected cells that express Esr1 is significantly higher than the percentage of total BNSTpr cells that express Esr1. n = 3 mice. Error bars: SEM. Unpaired T-test. ***p < 0.001. **(m-t)** Optogenetic activation of BNSTpr^Esr1^ cells in virgin and lactating C57BL/6 females induces immediate infanticide. Figure conventions as in **a-h**. n = 6 mice for GFP control, n = 8 mice for ChR2 virgin group, n = 6 mice for ChR2 mother group. Source data provided. Details of the statistical analyses and sample sizes can be found in Statistic Summary Table.

To understand whether BNSTpr^MPOA^ drive attack specifically towards pups or any social stimuli, we activated BNSTpr^MPOA^ in the presence of an adult female or male intruder (**Extended Data Figs. 3a-3h**). Only 1/10 tested females attacked adult intruders and only in < 3% of total stimulation trials (2/361 trials towards female intruders, 10/360 trials towards male intruders), suggesting that BNSTpr^MPOA^ does not drive attack towards adults consistently (**Extended Data Figs. 3a-3c and 3e-3g**). Besides attack, half of the animals showed male-style mounting occasionally (25/361 trials towards female intruders, 18/360 trials towards male intruders) (**Extended Data Figs. 3a-3c and 3e-3g**). Interestingly, all animals (10/10 towards female intruders, 8/10 towards male intruders) showed social grooming in approximately half of the total trials (208/361 trials towards female intruders, 135/360 trials towards male intruders) with an average latency of 3s (**Extended Data Figs. 3a-3h**).

Stress can negatively impact pup caring behaviors in mothers^43,44^ and several BNST subdivisions (though not BNSTpr) have been shown to modulate stress and anxiety^45–47^. To understand whether BNSTpr^MPOA^ stimulation evoked infanticide is due to an increase in anxiety, we examined light evoked behavioral changes in a real-time place preference test (RTPP) and an elevated plus maze test (EPM) (**Extended Data Figs. 3i and 3j**). We found that activation of BNSTpr^MPOA^ increased the amount of time spent on the stimulated side (**Extended Data Fig. 3i**) and open arms (**Extended Data Fig. 3j**), suggesting that BNSTpr^MPOA^ stimulation is not aversive or anxiogenic. Thus, the stimulation-induced infanticide is not secondary to an increase in stress.

### Optogenetic activation of BNSTpr^Esr1^ cells drives unconditional and immediate infanticide

Esr1 is widely expressed in regions important for social behaviors^48^. Previous studies showed that Esr1 positive (Esr1+) cells in VMHvl, MPOA and PA are preferentially involved in social behaviors in comparison to Esr1 negative (Esr1-) cells^49–51^. Within the BNST, Esr1 is concentrated in BNSTpr and expressed at a higher level in females than males^52^ (**Extended Data Fig. 3k**). Thus, we next investigated the possibility that Esr1 is a relevant molecular marker for infanticide cells in the BNST. Immunostaining revealed that Esr1 is expressed in approximately half of the BNSTpr cells and strikingly over 90% infanticide induced Fos+ cells overlap with Esr1+ cells (**Figs. 2i-2j**) and approximately 85% of MPOA-connecting BNSTpr cells are Esr1+ (**Figs. 2k-2l**). Thus, Esr1 preferentially marks the BNSTpr population activated by infanticide and largely encompasses the cells connected with MPOA.

We then optogenetically activated BNSTpr^Esr1^ neurons bilaterally by injecting Cre-dependent ChR2-EYFP virus into BNSTpr of virgin Esr1-2A-Cre C57BL/6 female mice (**Figs. 2m-2o**). Control animals were injected with a Cre-dependent GFP virus. We found that optogenetic activation of BNSTpr^Esr1^ cells induced pup attack even more reliably and quickly than BNST^MPOA^ cell activation (**Figs. 2p-2q**). Regardless of the light intensity, infanticide was induced in all tested females (8/8) and in 92% of total trials (33 to 89 trials per animal) with an average latency of 1s (**Figs. 2r-2t**). The expression of infanticide is perfectly locked to light stimulation. None of the test animals showed spontaneous pup attack without light and none of the GFP control animals attacked the pups during the entire test session (**Figs. 2p-2r**). We further asked whether BNSTpr^Esr1^ activation can override the maternal behaviors in mothers. Indeed, light stimulation induced reliable infanticide in all lactating females (6/6) even towards their own pups while all mothers quickly retrieved and cared for pups in the absence of light stimulation (**Figs. 2q-2t**).

We also examined BNSTpr^Esr1^ activation induced behavioral changes towards adult conspecifics (**Extended Data Figs. 3l-3s**). Similar to BNSTpr^MPOA^ activation, we observed a significant increase in social grooming towards both adult females (**Extended Data Figs. 3l-3o**) and adult males in nearly all tested animals (**Extended Data Figs. 3p-3s**). 50% (4/8) of females showed male-style mounting and attack in approximately 20% of total stimulation trials (**Extended Data Figs. 3l-3n and 3p-3r**).

Lastly, to ensure that the function of BNSTpr^Esr1^ cells is not specific to C57BL/6 females, we carried out the optogenetic activation in non-infanticidal Esr1-2A-Cre females with SW background and observed similar results in both virgin and lactating females (**Extended Data Figs. 3t-3w**). Collectively, these results indicate that BNSTpr^Esr1^ cells are sufficient to drive infanticide in females regardless of animals’ reproductive state and genetic background.

### BNSTpr^Esr1^ neurons are required for infanticide in females

To determine whether BNSTpr^Esr1^ neurons are necessary for infanticide in female mice, we bilaterally expressed hM4Di-mCherry in BNSTpr^Esr1^ cells in virgin Esr1-2A-Cre SW mice and tested their behavior towards pup in virgin and lactating states 30 minutes after saline or CNO injection (**Figs. 3a-3c**). Mice with SW background, instead of C57BL/6 background, were used given the higher rate of spontaneous infanticide of SW virgin females. Control mice expressed mCherry in BNSTpr^Esr1^ cells and were of SW background. CNO injection in control mice did not change the percentage of females that show infanticide or any other pup-directed behaviors (**Figs. 3d-3k**). In contrast, CNO injection in spontaneously infanticidal hM4Di-mCherry animals significantly reduced the fraction of females that killed pups (**Figs. 3d-3f**). Specifically, only 1/9 (11%) tested females attacked pups after CNO injection, whereas all 9 females killed pups after saline injection during the 10 minutes pup interaction test (**Figs. 3e-3f**). Furthermore, the fraction of females that showed pup retrieval significantly increased from 0/9 (0%) to 6/9 (66.7%) after CNO injection, supporting the hypothesis that infanticidal circuit and maternal circuit counteract one another and suppression of one circuit facilitates activation of the other (**Figs. 3e and 3g**). In females that did not show infanticide, CNO injection did not cause any change in pup directed behaviors, including the percentage of animals that retrieved pups, the latency to pup retrieval and duration of pup investigation **(Figs. 3h-3k**). Similarly, inactivation of BNSTpr^Esr1^ cells in lactating females did not change any measurements of pup-directed behaviors (**Figs. 3l-3o**). All lactating females showed reliable pup retrieval after saline or CNO injection (**Figs. 3l-3o**). Maternal aggression towards adult male and female intruder was also unaffected by BNSTpr^Esr1^ cell inactivation (**Extended Data Fig. 4**). All tested lactating females (11/11) attacked the intruder after both saline and CNO injection. The latency to attack (**Extended Data Figs. 4c and 4h**), the duration of attack (**Extended Data Figs. 4d and 4i**), and the total number of attacks (**Extended Data Figs. 4e and 4j**) are comparable after saline or CNO injection. Thus, inactivation of BNSTpr^Esr1^ cells in females specifically suppressed infanticide without changing any other aspects of pup-directed behaviors or aggressive behaviors towards adults.

**Fig. 3:**
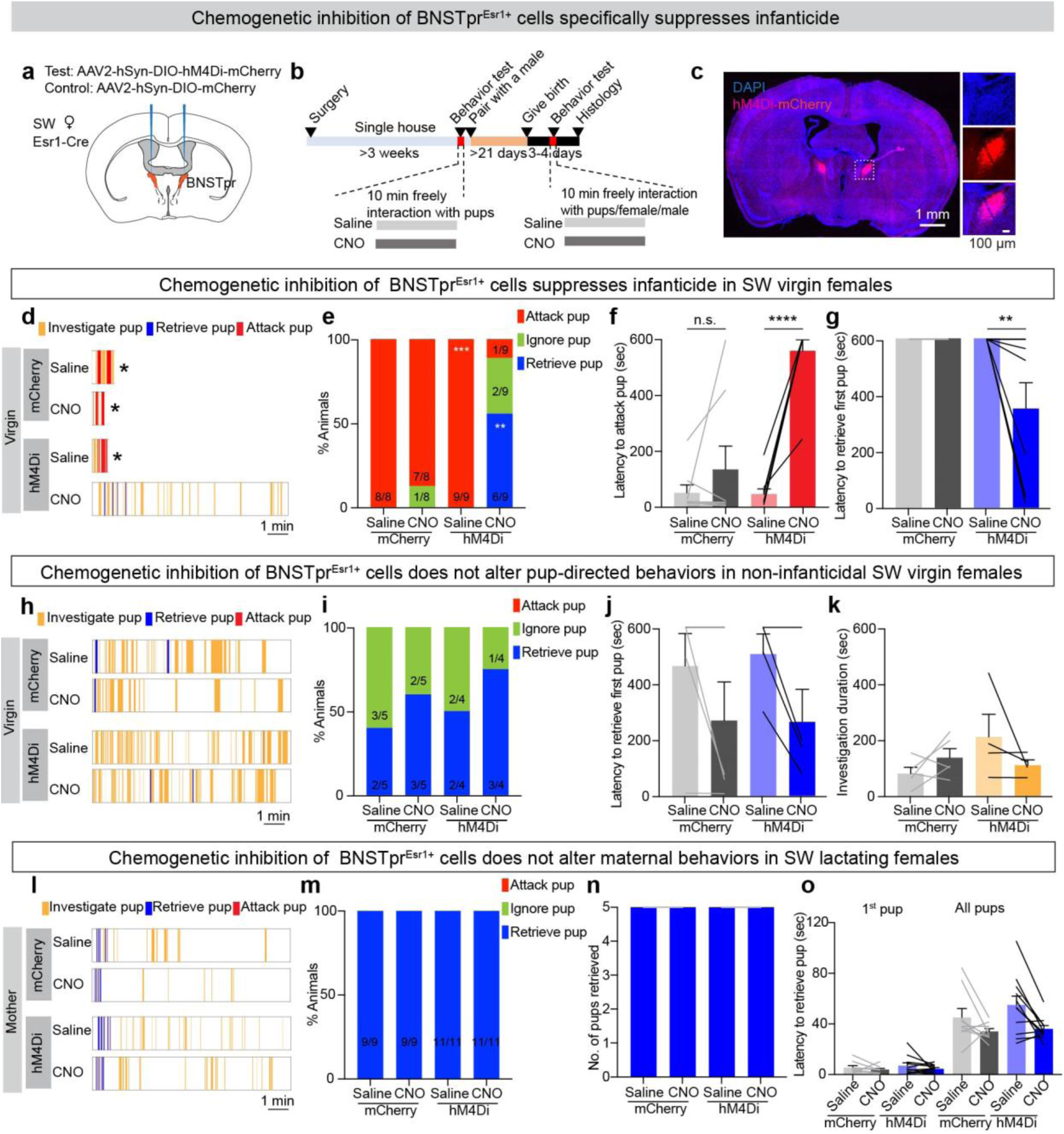
BNSTpr^Esr1^ neurons are required for infanticide in virgin females. **(a)** Experimental design to chemogenetically inhibit BNSTpr^Esr1^ neurons in SW females. **(b)** Experimental timeline. **(c)** A representative histology image showing hM4Di-mCherry (red) expression in BNSTpr region. **(d)** Representative raster plots showing pup-directed behaviors in mCherry control and hM4Di test females after i.p. injection of CNO or saline. *Remove hurt pups and stop recording. **(e)** Percentage of control and test virgin females that attack, ignore and retrieve pups after saline and CNO injection. All females included here attacked pups spontaneously. Fisher’s exact test. **p < 0.01. ***p < 0.001. **(f, g)** Latency to attack pup (**f**) or retrieve pup (**g**) after saline or CNO injection in hostile virgin control and test females. Error bars: SEM. Two-way ANOVA with Bonferroni’s multiple comparisons test. **p < 0.01, ****p < 0.0001. n = 8 for mice mCherry control, n = 9 mice for hM4Di group. **(h)** Representative raster plots showing pup-directed behaviors in non-hostile mCherry and hM4Di virgin females after saline or CNO injection. **(i)** Percentage of control and test virgin females that attack, ignore and retrieve pups after saline or CNO injection. None of the females were spontaneously infanticidal. **(j-k)** Latency to retrieve pup (**j**) and pup investigation duration (**k**) after saline or CNO injection in non-hostile virgin control and test females. Error bars: SEM. n = 5 for mice mCherry control, n = 4 mice for hM4Di group. **(l)** Representative raster plots showing various pup-directed behaviors in lactating mCherry and hM4Di females after saline or CNO injection. **(m)** 100% of control and test lactating females retrieved pups after saline or CNO injection. **(n)** All 5 pups were retrieved in 10 minutes testing period in control and test lactating females after either saline or CNO injection. n = 9 for mice mCherry control, n = 11 mice for hM4Di group. **(o)** Latency to retrieve the first pup and all five pups in control and test lactating females after either saline or CNO injection. Error bars: SEM. n = 9 for mice mCherry control, n = 11 mice for hM4Di group. Source data provided. Details of the statistical analyses and sample sizes can be found in Statistic Summary Table.

### BNSTpr^Esr1^ and MPOA^Esr1^ cells show opposite response pattern during maternal care and infanticide in females

Our functional manipulations demonstrated that BNSTpr^Esr1^ neurons are sufficient and necessary for infanticide in female mice. To understand the responses of BNSTpr^Esr1^ cells under natural pup-directed behaviors, we utilized fiber photometry to record population activity of BNSTpr^Esr1^ cells in Esr1-2A-Cre SW females (**Figs. 4a-4b**). Esr1 immunostaining indicated that over 90% GCaMP6f+ cells express Esr1, confirming that BNSTpr^Esr1^ cells are the main source of the fluorescence signal (**Fig. 4c**). Each animal was recorded in hostile virgin and lactating state to understand the cell responses during various pup-directed behaviors (**Figs. 4f-4m**).

**Fig. 4:**
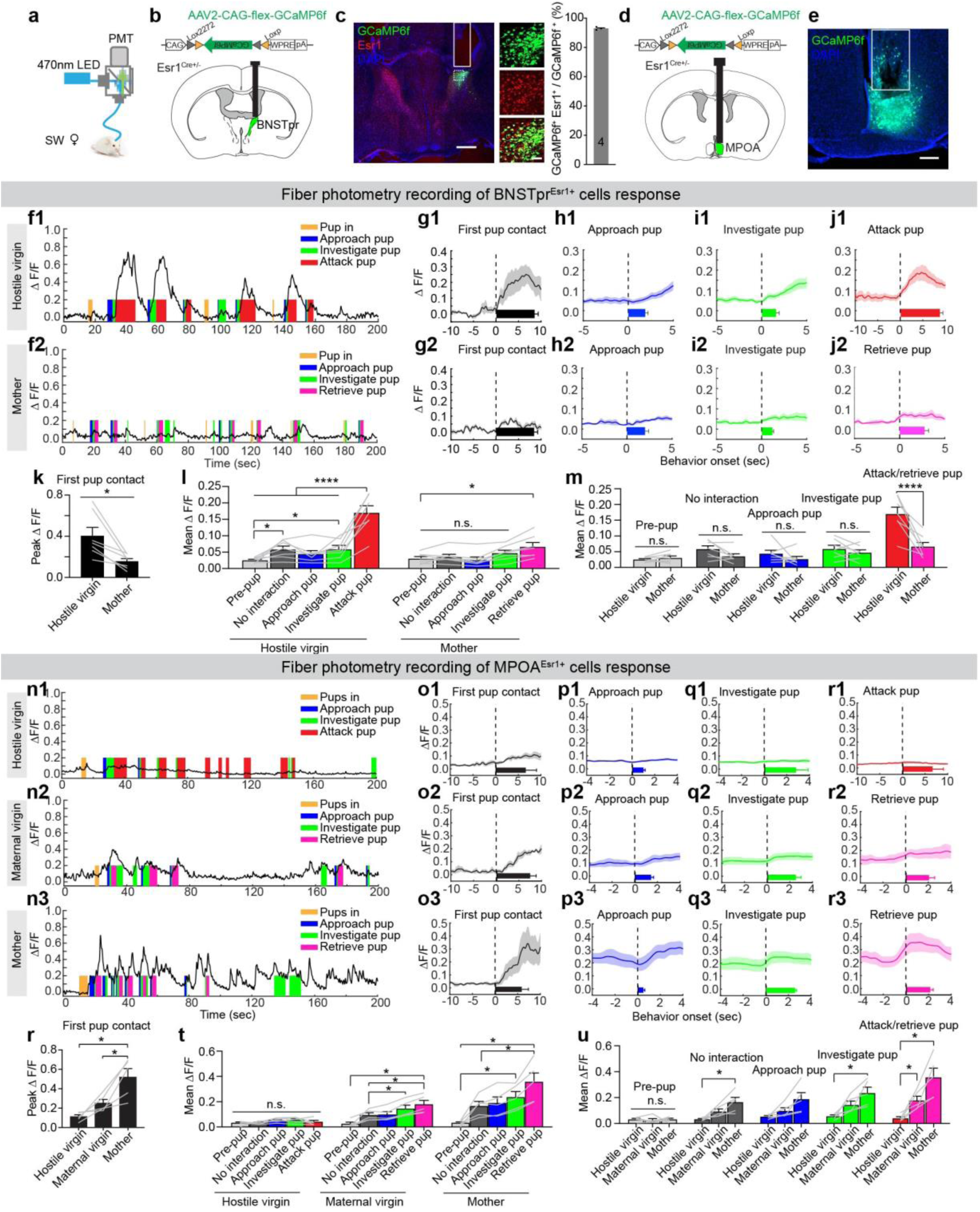
Opposite responses of BNSTpr^Esr1^ and MPOA^Esr1^ cells during maternal care and infanticide in females. **(a)** Fiber photometry setup. (**b and d**) viral construct and targeted brain regions. **(c)** A representative histological image showing the fiber track in BNSTpr (white line), the overlap between Esr1 staining (red) and GCaMP6f (green). The enlarged view of the boxed area is shown to the right. Scale bars: 500 µm (left) and 50 µm (right). Bar graph shows the averaged percentage of the GCaMP6f cells that express Esr1 from 4 recording mice. Error bars: SEM. **(d)** A representative histological image showing the expression of GCaMP6f (green) and fiber track (white line) in the MPOA (**e**). Blue, DAPI. Scale bar: 200 µm. **(f)** Representative GCaMP6f recording (ΔF/F) traces during pup interaction from a hostile virgin female (**f1**) and a mother (**f2**). Color shades indicate various behaviors. **(g-j)** Average post-event histograms (PETHs) of GCaMP6f signal (ΔF/F) aligned to the onset of various behaviors in hostile virgin females (**g1-j1**), and mothers (**g2-j2**). n = 7 mice. Horizontal bars in the graph indicate the average duration of a behavior. Shades and error bars: ± SEM. **(k)** Peak response (ΔF/F) during the first pup contact. Paired t-test. *p < 0.05, Error bars: SEM. **(l-m)** Mean GCaMP6f signal (ΔF/F) during various pup-directed behaviors and pre-pup introduction period in hostile virgin females and mothers. Two-way. *p < 0.05, ****p < 0.0001, Error bars: SEM. **(n)** Representative GCaMP6f recording (ΔF/F) traces during pup interaction from a hostile virgin female (**n1**), a maternal virgin female (**n2**) and a mother (**n3**). Color shades indicate various behaviors. **(o-r)** PETHs of GCaMP6f signal (ΔF/F) aligned to the onset of various behaviors towards pups in hostile virgin females (**o1-r1**), maternal virgin females (**o2-r2**) and mothers (**o3-r3**). n = 5 mice. Horizontal bars indicate the average duration of the behavior events. Shading and error bars: SEM. **(s)** Peak responses (ΔF/F) during first pup contact. One-way ANOVA followed with Tukey’s multiple comparison test. *p < 0.05, Error bars: SEM. **(t-u)** Mean GCaMP6f signal (ΔF/F) during various pup-directed behaviors and pre-pup introduction period in hostile virgin females, maternal virgin females and mothers. Two-way ANOVA followed with Tukey’s multiple comparison test. *p < 0.05, Error bars: SEM. Source data provided. Details of the statistical analyses and sample sizes can be found in Statistic Summary Table.

During the first pup contact after its introduction, we observed a sharp rise of Ca^2+^ signal in hostile virgin females, but not in lactating females (**Figs. 4f, 4g and 4k)**. For each subsequent episode of interaction, Ca^2+^ signal did not significantly rise during pup approach and increased only slightly during close pup investigation (**Figs. 4h, 4i and 4l**). When the hostile female initiate attack towards a pup, Ca^2+^ signal rose sharply and maintained at the high level until the end of attack (**Fig. 4j1**). In contrast, BNSTpr^Esr1^ cells only increased activity slightly during pup retrieval in mothers (**Figs. 4j2 and 4l)**. The average response of BNSTpr^Esr1^ cells during infanticide is significantly higher than that during retrieval (**Fig. 4m**). When females did not actively interact with pups, GCaMP6f signal did not change from the pre-pup level in mothers but increased slightly in hostile virgin females (**Fig. 4l**). Overall, BNSTpr^Esr1^ cells show large activity increase specifically during infanticide (**Figs. 4l and 4m**).

We next asked whether BNSTpr^Esr1^ cells respond during attack towards adult conspecifics by introducing a male or a female intruder into the home cage of the recording lactating female (**Extended Data Fig. 5a**). When the recording mouse first encountered an adult intruder, we observed a large increase in Ca^2+^ signal (**Extended Data Figs. 5a, 5b and 5e**). During subsequent male or female investigation, there is no clear response (**Extended Data Figs. 5c and 5f**). Importantly, during attack towards an adult intruder, there is little activity increase of BNSTpr^Esr1^ cells (**Extended Data Fig. 5d**). Overall, the response of BNSTpr^Esr1^ cells is significantly higher during pup attack than adult attack (**Extended Data Fig. 5g**).

The responses of BNSTpr^MPOA^ cells during pup and adult interactions were qualitatively similar to those of BNSTpr^Esr1^ cells. Interestingly, BNSTpr^MPOA^ cells showed overall higher responses to pups in hostile females and lower responses to pups in lactating females in comparison to BNSTpr^Esr1^ cells, suggesting that infanticide-responsive BNSTpr cells, but not retrieval-responsive cells, preferentially project to MPOA (**Extended Data Figs. 5h-5v**). We further recorded Ca^2+^ signal of MPOA^Esr1^ cells during pup interaction in Esr1-2A-Cre SW females before and after the animals changed from infanticidal to maternal with pup exposure and parturition (**Figs. 4d-4e**). The responses of MPOA^Esr1^ cells to pups were the direct opposite to those of BNSTpr^Esr1^ cells (**Figs. 4n-4u**). In virgin hostile females, MPOA^Esr1^ cells activity increased minimally during approach and investigation and not at all during attack pups (**Figs. 4n1-4r1 and 4t**). In maternal virgin females and mothers, Ca^2+^ signal started to rise when the female approached the pup, continued to increase during investigation and reached maximum during retrieval (**Figs. 4n2-4q2, 4n3-4q3 and 4t**). During retrieval, the average fluorescence increase is higher in mothers than that in virgin females (**Figs. 4r2**, **4r3 and 4u**). We also noticed a higher baseline Ca^2+^ activity in mothers than in virgin infanticidal females during the period when the female did not interact with the pups (**Fig. 4u**). Thus, the activity of MPOA^Esr1^ cells vary with maternal state and increases during positive but not negative pup directed behaviors.

MPOA^Esr1^ cells also increase activity upon introduction of an adult intruder and during each subsequent interaction (**Extended Data Figs. 6a-6j**). However, unlike responses to pups, the average responses during investigation of adult conspecific are similar in hostile virgin females, maternal virgin females and mothers (**Extended Data Figs. 6e and 6j**). When comparing among the stimuli, MPOA^Esr1^ cells respond more to adult conspecifics than pups in hostile females while the opposite is true in mothers (**Extended Data Figs. 6k and 6l**).

Altogether, consistent with the functional results showing an indispensable role of BNSTpr^Esr1^ cells in infanticide, the cells are activated strongly during infanticide and minimally during maternal behaviors, which is directly opposite to the response pattern of MPOA^Esr1^ cells.

### BNSTpr^Esr1^ and MPOA^Esr1^ neurons form reciprocal inhibition

The opposing response pattern of BNSTpr^Esr1^ and MPOA^Esr1^ neurons is consistent with the hypothesis that these two populations counteract one another. We next examined the axon terminals of BNSTpr^Esr1^ and MPOA^Esr1^ cells by virally expressing GFP in those cells. We found dense terminal fields in Esr1-enriched region in the MPOA originated from BNSTpr^Esr1^ cells and vice versa (**Extended Data Figs. 7a-7c and 7f-7h**). A survey of the terminal fields throughout the brain revealed that MPOA represents one of the major downstream regions of the BNSTpr^Esr1^ whereas BNSTpr receives moderate input from MPOA^Esr1^ cells (**Extended Data Figs. 7d-7e and 7i-7j)**.

To address whether BNSTpr^Esr1^ cells provide direct synaptic input to MPOA^Esr1^ cells, we employed ChR2-assisted circuit mapping by injecting Cre-dependent ChR2-EYFP into BNSTpr and Cre-dependent mCherry into MPOA of SW Esr1-2A-Cre virgin female mice (**Figs. 5a and 5b**). The viruses were reversed for probing MPOA^Esr1^ to the BNSTpr^Esr1^ projection (**Figs. 5f and 5g**). Three weeks later, we performed whole-cell voltage clamp recordings from mCherry+ cells in MPOA or BNSTpr on slices while stimulating BNSTpr^Esr1^ or MPOA^Esr1^ terminals using 0.5-ms blue-light pulses. Vast majority of MPOA^Esr1^ neurons (91%, 20/22) showed optogenetically evoked inhibitory postsynaptic currents (oIPSCs) including two neurons showing both oIPSCs and evoked excitatory postsynaptic currents (oEPSCs) (**Fig. 5c**). The oIPSC was large (Mean ± SEM: 930 ± 166 pA) and monosynaptic as it was blocked by tetrodotoxin (TTX) and rescued by bath application of a mix of TTX and 4-aminopyridine (4-AP) (**Figs. 5d-5e**). The oIPSC is mediated mainly by GABA_A_ receptors and to a lesser extent by glycine receptors: GABA_A_ receptors antagonist, gabazine (SR), completely blocked oIPSC in 56% (20/36) of cells (**Figs. 5d-5e and Extended Data Figs. 7k-7l**) and the residual oIPSCs can be further blocked by glycine receptor inhibitor Strychnine (**Extended Data Figs. 7m and 7n**).

**Fig. 5:**
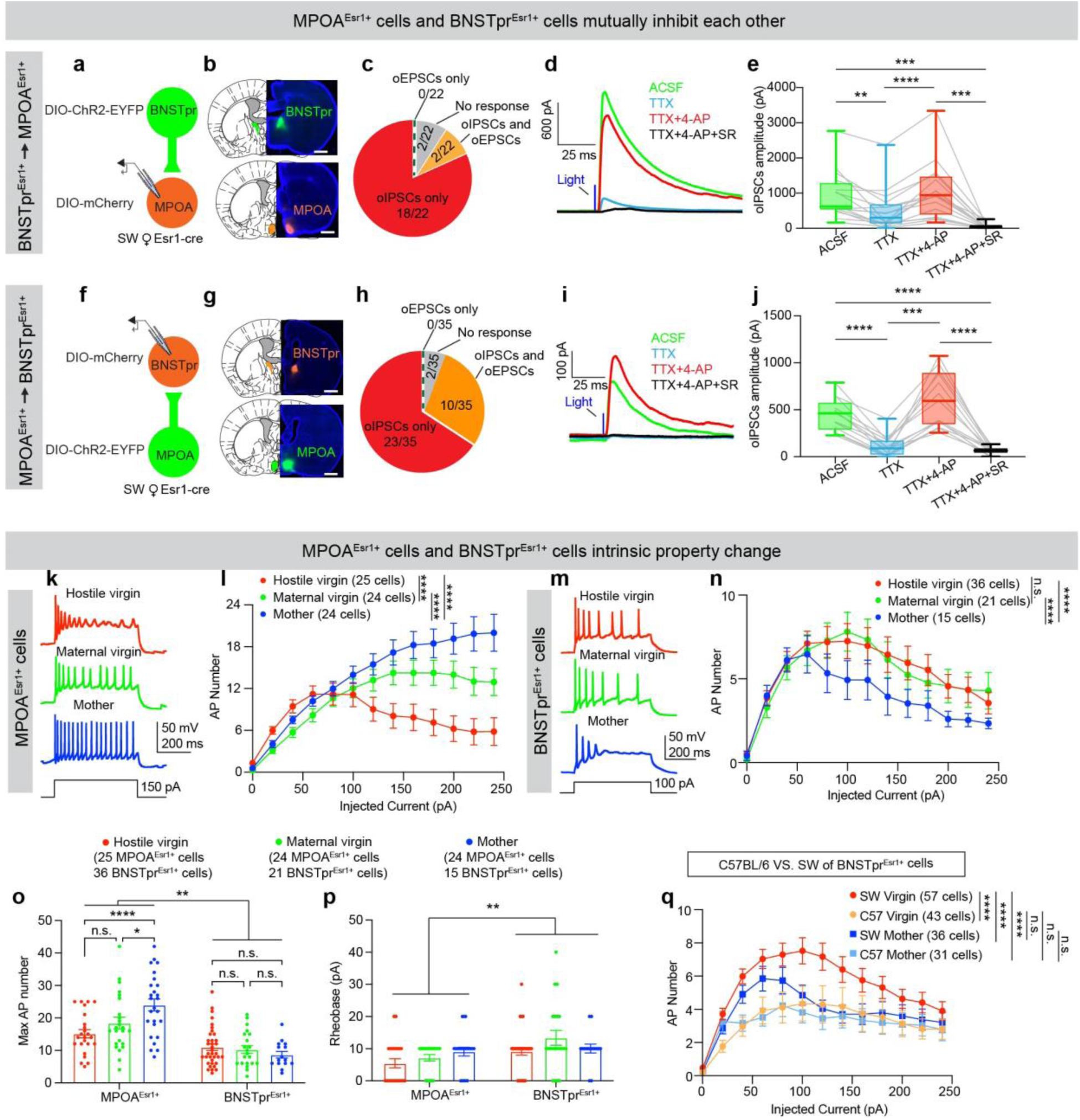
Mutual inhibition of BNSTpr^Esr1^ and MPOA^Esr1^ cells, and their Intrinsic property change. **(a and f)** Schematics of ChR2-assisted circuit mapping of BNSTpr^Esr1^ → MPOA^Esr1^ (**a**) and MPOA^Esr1^ → BNSTpr^Esr1^ (**f**). **(b and g)** Representative images showing ChR2-EYFP in BNSTpr^Esr1^ and mCherry in MPOA^Esr1^ (**b**), and ChR2-EYFP in MPOA^Esr1^ and mCherry in BNSTpr^Esr1^ (**g**). Blue: DAPI. Scale bars: 1 mm. **(c and h)** Pie charts showing the synaptic responses of MPOA^Esr1^ cells to BNSTpr^Esr1^ terminal activation **(c)** and responses of BNSTpr^Esr1^ cells to MPOA^Esr1^ terminal activation (**h**). n = 22 cells from 3 mice in (**c**) and 35 cells from 3 mice in (**h**). **(d and i)** Representative oIPSC traces from MPOA^Esr1^ cells (**d**) and BNSTpr^Esr1^ cells (**i**) with different blockers. **(e and j)** oIPSC amplitude with different blockers. One-way ANOVA followed with Tukey’s multiple comparison test. **p < 0.01, ***p < 0.001, ****p < 0.0001, Error bars: min to max. n = 16 cells from 4 mice in (**e**) and 13 cells from 5 mice in (**j**). **(k)** Representative recording traces of MPOA^Esr1^ cells from a hostile virgin female (top), a maternal virgin female (middle) and a mother (down) with 150 pA current injection. **(l)** F-I curves of MPOA^Esr1^ cells from hostile virgin females (red), maternal virgin females (green) and mothers (blue). Two-way ANOVA. ****p < 0.0001. Error bars: ± SEM. Each group contains 24-25 cells collected from 3 animals. **(m)** Representative recording traces of BNSTpr^Esr1^ cells from a hostile virgin female (top), a maternal virgin female (middle) and a mother (down) with 100 pA current injection. **(n)** F-I curves of BNSTpr^Esr1^ cells from hostile virgin females (red), maternal virgin females (green) and mothers (blue). Two-way ANOVA. ****p < 0.0001. Error bars: ± SEM. Each group contains 15 to 36 cells collected from 3 animals. **(o)** Maximum AP number of MPOA^Esr1^ and BNSTpr^Esr1^ cells from hostile virgin females (red), maternal virgin females (green) and mothers (blue) with current injections ranging from 0 pA to 250 pA. Two-way ANOVA. *p < 0.05, **p < 0.01, ****p < 0.0001. Error bars: ± SEM. **(p)** Rheobase of MPOA^Esr1^ and BNSTpr^Esr1^ cells from hostile virgin females (red), maternal virgin females (green) and mothers (blue). Two-way ANOVA. **p < 0.01. Error bars: ± SEM. Each group contains 15 to 36 cells from 3 animals. **(q)** F-I curves of BNSTpr^Esr1^ cells from C57BL/6 and SW female mice. Two-way ANOVA. ****p < 0.0001. Error bars: ± SEM. Each group contains 31 to 57 cells collected from 3-6 animals. Source data provided. Details of the statistical analyses and sample sizes can be found in Statistic Summary Table.

Similarly, 94% (33/35) BNSTpr^Esr1^ showed oIPSCs upon MPOA^Esr1^ terminal stimulation including 10 cells that showed both oIPSCs and oEPSCs (**Fig. 5h**). The higher proportion of cells showing oEPSCs upon MPOA^Esr1^ BNSTpr^Esr1^ stimulation in comparison to BNSTpr^Esr1^ MPOA^Esr1^ stimulation is consistent with the fact that BNSTpr^Esr1^ cells are nearly exclusively GABAergic, whereas approximately a quarter of MPOA^Esr1^ cells are glutamatergic^26,53^. The oIPSCs can be blocked by bath application of TTX and rescued by TTX and 4-AP, suggesting the monosynaptic nature of the connection (**Figs. 5i and 5j**). The oIPSCs are largely mediated by GABA_A_ receptor and to a smaller extent by glycine receptor (**Figs. 5i-5j and Extended Data Figs. 7o-7r**). Consistent with the difference in terminal field density, the monosynaptic oIPSC from BNSTpr^Esr1^ MPOA^Esr1^ is higher than that of MPOA^Esr1^ BNSTpr^Esr1^ (**Extended Data Figs. 7s and 7t**). Together, these results suggested that BNSTpr^Esr1^ and MPOA^Esr1^ neurons form strong reciprocal inhibitory connections and likely could counteract each other.

### The excitability of BNSTpr^Esr1^ and MPOA^Esr1^ neurons change with reproductive states

After parturition, all females showed robust maternal behaviors even if they were infanticidal during the virgin state. We next asked whether this change in pup-directed behavior is supported by a shift in balance between MPOA^Esr1^ and BNSTpr^Esr1^ cell activity. We performed *in vitro* current clamp recording of MPOA^Esr1^ and BNSTpr^Esr1^ cells from diestrus hostile virgin, diestrus maternal virgin and lactating (postpartum day 3) SW Esr1-2A-Cre female mice. For each animal, we recorded both MPOA^Esr1^ and BNSTpr^Esr1^ cells to eliminate the contribution of individual variability within a reproductive state. We found distinct state-dependent changes in MPOA^Esr1^ and BNSTpr^Esr1^ cell excitability. MPOA^Esr1^ cells in hostile virgin females were prone to depolarization block and did not maintain high spiking activity with moderate level of current injection (>100pA), whereas MPOA^Esr1^ cells in mothers continued to increase firing with current injection (**Figs. 5k and 5l**). The excitability of MPOA^Esr1^ cells in maternal virgins are in between that of hostile virgins and mothers (**Figs. 5k and 5l**). In contrast, BNSTpr^Esr1^ cells in mothers were less excitable than those in virgin females as reveled by the spike frequency current (F-I) curves, although the excitability of BNSTpr^Esr1^ cells in hostile and maternal virgin females were similar (**Figs. 5m and 5n**). Between BNSTpr^Esr1^ and MPOA^Esr1^ cells, MPOA^Esr1^ were generally more active as reflected by their higher maximum action potential (AP) number during current injection and lower rheobase, and this difference was the largest in mothers (**Figs. 5o and 5p**). Overall, MPOA^Esr1^ cells are more excitable in mothers than virgin females, while the opposite is true for BNSTpr^Esr1^ cells. These opposing changes in excitability of MPOA^Esr1^ and BNSTpr^Esr1^ cells could be essential for the quick emergence of maternal behaviors in mothers.

Lastly, we asked whether the difference in the tendency to express infanticidal behaviors between SW and C57BL/6 virgin females could reside in their differences in BNSTpr^Esr1^ cell properties. We performed current clamp recording of BNSTpr^Esr1^ cells from virgin C57BL/6 females (BNSTpr^Esr1.C57)^ and found that they were much less excitable than those in SW females (BNSTpr^Esr1.SW^) (**Fig. 5q**). In fact, approximately half of BNSTpr^Esr1.C57^ cells (23/44) could not be driven to fire more than two spikes regardless of the amount of injected current, whereas the same was true for only 1/57 BNSTpr^Esr1.SW^ cells in virgin females (**Extended Data Figs. 7u-7v**). Across the populations, the spiking frequency of BNSTpr^Esr1.C57^ cells was significantly lower than that of BNSTpr^Esr1.SW^ cells in virgin females at all current steps while the F-I curves of BNSTpr^Esr1.C57^ and BNSTpr^Esr1.SW^ cells in lactating animals did not differ (**Fig. 5q**). These results revealed dampened excitability of BNSTpr^Esr1^ cells in virgin C57BL/6 females, which likely contribute to a lack of infanticidal behaviors in these animals.

## Discussion

Though initially considered as a pathological behaviors shown only in rare occasions, infanticide may in fact be an adaptive behaviour to increase an inidividual’s reproductive success in both males and females^13,15^. Recent studies started to reveal a circuit dedicated for infanticide in males, but the existence of such a circuit in females and its identity remains elusive. Here, using MPOA as an entry point, we identified three regions, BNSTpr, MeApd and VMHvl, that can drive negative pup-directed behaviors in female mice, each with differential behavioral outputs. We further investigated the BNSTpr^Esr1^ population in detail and established its indispensable role in female infanticide and its close interaction with MPOA^Esr1^ cells in controlling maternal behavior (**Extended Data Fig. 8**). The relative activities between BNSTpr^Esr1^ and MPOA^Esr1^ cells shift with reproductive states, which could be responsible for the dramatic changes in pup-directed behaviors during motherhood.

### The role of BNSTpr in infanticide and other social behaviors

BNSTpr has long been suggested as a part of the social behavior network given its abundant expression of sex steroid receptors, its strong connection with medial hypothalamic regions and its direct input from that accessory olfactory bulb (AOB) that carries pheromone cues of conspecifics^54^. BNSTpr is a sexually dimorphic region with a larger size in males than females and thus most studies have been carried out in males^55,56^. In males, early lesions and recent functional studies on BNSTpr aromatase or CCK expressing cells suggest a role of the region in mediating innate sex preference and promoting copulation when the animals are sexually inexperienced whereas the BNSTpr function in females are largely unknown^57–65^. Female BNSTpr aromatase expressing cells do not respond to any tested social cues (male, female and pup in mothers) and is unnecessary for sex recognition, mating or maternal aggression^65^. Although high level of c-Fos is observed in BNSTpr after female sexual behaviors^66^, pelvic nerve transection eliminates the BNSTpr c-Fos without impairing female sexual behaviors, which argues against an important role of the cells in the expression of female sexual behaviors^67^.

Here, we found that BNSTpr^Esr1^ are necessary, sufficient, and highly activated during infanticide in female mice, assigning a clear social function to the female BNSTpr cells. Unexpectedly, although BNSTpr receives direct AOB input that presumably carry information of pheromones, the responses of BNSTpr^Esr1^/BNSTpr^MPOA^ cells during pup investigation are quite modest in comparison to those during infanticide. The strong action associated BNSTpr^Esr1^ cell responses are consistent with the short latency (<1s) to initiate pup attack upon optogenetic stimulation of the cells. The response of BNSTpr^Esr1^ cells during attacking pups is significantly higher than that during attacking adult intruders. This response pattern contrasts with that of posterior substantia innominate (pSI), where cells respond more strongly when attacking adult intruders as compared to pups^68^. Consistent with the pup-specific response pattern of BNSTpr^Esr1^ cells, activating the cells at the lowest light intensity (0.5 mW) drives infanticide with near 100% efficiency while stimulating the cells using the maximum light (> 5mW) failed to induce attack towards adult intruders consistently. In contrast, activating pSI drives attack towards nearly all social and non-social targets^68^. When BNSTpr^Esr1^ cells are inactivated, infanticide is nearly abolished while maternal aggression is unaffected. Thus, BNSTpr^Esr1^ cells mediate aggression specifically towards pups. This result suggests that infanticide and adult aggression are mediated by differential neural circuits even though they may converge at certain points, e.g., pSI^34,68,69^.

When BNSTpr^Esr1^ cells were artificially activated, it also induced social grooming towards both adult male and female intruders in approximately half of the trials. It is unclear whether the induced social grooming represents an affiliative behavior or a low-intensity agonistic behavior. During our *in vivo* recording, we rarely observed spontaneous social grooming, precluding our ability to investigate the endogenous cell responses during this behavior. Interestingly, Wu et. al. recently reported social grooming induced by optogenetic activation of MeA GABAergic cells that express tachykinin (MeA^Tac1⋂Vgat^) and suggested that social grooming mainly serves a role in comforting stressed conspecifics^70^. Given the strong connection between MeApd and BNSTpr, it is possible that MeA^TaC1⋂Vgat^ and BNSTpr^Esr1^ cells activate the same circuit to drive social grooming. Future studies will elucidate the importance of BNSTpr^Esr1^ cells in naturally occurring social grooming and the relationship between infanticide- and social grooming-responsive BNSTpr^Esr1^ cells at the single cell level.

### Infanticide circuit in females beyond BNSTpr

Anosmia and lesion in the MeA or medial hypothalamus facilitate the emergence of maternal behaviors in virgin female rats^71–75^. These functional results along with the higher immediate early gene expression in these areas with pup exposure in non-maternal females compared to maternal females, led to the hypothesis of a maternal behavior-suppression circuit driven by pup odor^6,76^. These lesion studies, however, did not directly address whether any specific pup-directed behaviors are mediated by this suppression circuit. Here, using systematic gain of function manipulation, we found that in addition to BNSTpr^MPOA^ cells, MeA^MPOA^ cells drive pup grooming and infanticide, while VMHvl^MPOA^ cells promote pup avoidance.

A previous study found that optogenetic activation of MeA GABAergic cells induce pup grooming but not infanticide in female mice^34^. The lack of stimulation-induced infanticide in the previous study might suggest that MeA^MPOA^ cells are more specifically relevant for infanticide than the GABAergic cells, which constitute the major population in the MeApd^77^. We are uncertain whether the MeA^MPOA^ stimulation-induced pup grooming represents a nurturing maternal behavior or motivated by aggression since we observed pup grooming in both infanticidal and maternal females naturally. Future recording from MeA cells during pup grooming in animals under different maternal states will help address this question. MeA and BNSTpr both receive direct inputs from the AOB and are strongly reciprocally connected. Thus, we speculate that MeA and BNSTpr represents two nodes in the same circuit. Consistent with this hypothesis, MeA lesion reduced the pup induced c-Fos in the unilateral BNSTpr in non-maternal female rats^78^.

Despite a well-known role of VMHvl in adult-directed aggression, activating VMHvl^MPOA^ cells did not elicit attack towards pups^79^. Instead, the test female approached the pup repeatedly but then quickly retreated after a brief interaction a characteristic avoidant behavior. Thus, although both BNSTpr and MeA project to VMHvl, it is unlikely that BNSTpr and MeA mediate infanticide through VMHvl. Instead, VMHvl^MPOA^ may be a part of the circuit driving pup avoidance. Indeed, VMHvl is known to contain a population of cells relevant for avoidance towards social threats^80–82^. These results also suggest that infanticide and pup avoidance could be mediated by distinct neural circuits.

Lateral to the BNSTpr is the rhomboid nucleus of BNST (BNSTrh), a region that has been indicated in infanticide in male mice^33^. Consistent with the c-Fos pattern in males, we observed increased c-Fos expression in this area in virgin females after infanticide, indicating its potential relevance for female infanticide. However, BNSTrh is minimally connected with MPOA as we observed very few labeled cells in BNSTrh (< 10 cells/section vs. ∼175 cells/section in BNSTpr) traced from MPOA. Previous ChR2 assisted circuit mapping also failed to find a direct connection from MPOA to BNSTrh in males^33^. Additionally, Esr1 expresses only sparsely in this region (**Extended Data Fig. 3k**). Thus, our BNSTpr^MPOA^ and BNSTpr^Esr1^ manipulations are not expected to engage BNSTrh cells in any significant way. Does the BNSTpr/MeA and BNSTrh belong to the same infanticide circuit? Answer to this question remain unclear as classical tracing studies and our anterograde tracing from BNSTpr^Esr1^ cells failed to uncover any appreciable connections between BNSTrh and BNSTpr/MeA^83,84^. Whether these areas interact or form parallel pathways to drive infanticide remain to be investigated in future studies.

### Neural plasticity of maternal and infanticide circuits during motherhood

Motherhood is associated with a drastic increase in infant caring behaviors that is presumably supported by changes in the maternal circuit. Since the identification of MPOA as a key site for maternal behaviors, numerous studies have focused on understanding its plasticity during motherhood and revealed various changes in cell morphology and gene expression patterns. For example, Keyser Marcus et. al. found an increase in dendritic length and branching complexity of MPOA cells in postpartum female rats in comparison to virgin females^85^. Uriarte et. al. found that perineurial nets form around the MPOA^Esr1^ cells prior to parturition and then gradually fades away through the postpartum period^86^. At the molecular level, a variety of neuropeptides and receptors in the MPOA were found to alter during postpartum, with some increasing by multiple folds^87,88^. Surprisingly, despite the extensive studies on the structural and molecular changes of MPOA cells during motherhood, to our best knowledge, changes in the electrophysiological properties of MPOA cells during motherhood have not been reported. Here, our results revealed a significant increase in intrinsic excitability of MPOA^Esr1^ during early postpartum (day 3) when the maternal motivation is the highest^89,90^. This change in excitability matches with the increased *in vivo* response to pups in mothers.

In virgin females, MPOA^Esr1^ cells are more prone to depolarization block when the females show infanticide, suggesting a limited output capacity of cells even when they receive a strong excitatory drive. This result is consistent with the blunted *in vivo* responses of MPOA^Esr1^ cells to pup cues in infanticidal virgin females, as presented in Fig. 4. A previous study showed that estradiol depletion induces depolarization block of MPOA neurotensin expressing cells in female mice^91^. However, in our study, both infanticidal and maternal virgin females are normal cycling diestrus females, arguing against a major role of sex hormones in causing the difference in intrinsic excitability. Indeed, maternal behaviors can be induced in virgin females with repeated pup exposure in the absence of sex hormones^92–94^. The pup experience-induced maternal behaviors has been linked to activation of cascades of intracellular signaling events that lead to chromatin remodeling and gene transcription, which is likely also responsible for changes in MPOA cell excitability in maternal virgins^94^.

The electrophysiological properties of BNSTpr cells are virtually unknown. Our results showed that BNSTpr^Esr1^ cells are generally less excitable than MPOA^Esr1^ cells, with most cells firing no more than 10 Hz at maximum. In comparison to MPOA^Esr1^ cells, changes in the excitability of BNSTpr^Esr1^ cells over reproductive state are less dramatic: BNSTpr^Esr1^ cells of lactating females are slightly less excitable than those in virgin females, whereas no difference in BNSTpr^Esr1^ cell excitability was found between infanticidal and maternal virgin females. These results suggest that MPOA^Esr1^ cells are likely to play a bigger role than BNSTpr^Esr1^ cells in shifting the balance between the outputs of maternal and infanticide circuits.

Although BNSTpr^Esr1^ cell excitability does not differ between infanticidal and maternal virgin females, we noticed generally lower excitability of BNSTpr^Esr1^ cells in C57BL/6 females in comparison to SW females. The maximum firing of BNSTpr^Esr1.B6^ cells is approximately 50% of that of BNSTpr^Esr1.SW^ cells. Half of BNSTpr^Esr1^ cells in C57 females never fire more than two spikes during current injection. This extraordinary resistance to firing of BNSTpr^Esr1.B6^ cells may explain a lack of infanticidal behavior in C57 female mice. These results also suggest that behavioral traits of animals with different genetic backgrounds could be encoded in the intrinsic properties of cells in the relevant circuits.

A negative circuit that counteracts the maternal circuit has long been suspected^5,6^. Here, our study unequivocally demonstrated the existence of an infanticide circuit in females and revealed its plasticity over the reproductive state and its variability among individuals with different propensity to kill. Our study further uncovered the intimate and antagonistic relationship between the infanticide circuit and maternal circuit, highlighting the importance of studying both circuits to understand infant-directed behaviors under normal and pathological conditions.

## Methods

### Mice

All animal experiments were performed in accordance with NIH guidelines and approved by the New York University medical school Institutional Animal Care and Use Committee (IACUC). Esr1-2A-Cre mice were provided by D.J. Anderson and are currently available from Jackson Laboratory (stock no. 017911), Ai6 mice were purchased from the Jackson Laboratory (stock no. 007906). They were backcrossed to either SW or C57 for at least five generations. WT SW mice were purchased from Taconic. WT C57BL/6 and Balb/c mice were purchased from Charles River, P1-P5 pups used for behavioral experiments were from our breeding colony. Mice were housed in 12 h light-dark cycle (10 p.m. 10 a.m. light), with food and water available ad libitum. All mice were group housed until adult. After surgery, mice were single housed unless they were paired with a male, and after they became obvious pregnant, they were single housed again until having a litter.

### Virus

AAV2-CAG-Flex-GCaMP6f was purchased from the University of Pennsylvania vector core. AAV2-hSyn-FLEX-GFP and AAV2-EF1a-DIO-ChR2-EYFP were purchased from the University of North Carolina vector core. AAV1-hSyn-Cre, AAV2-hSyn-DIO-mCherry, AAV2-hSyn-DIO-hM3Dq-mCherry and AAV2-hSyn-DIO-hM4Di-mCherry were purchased from Addgene. The titer of AAV1-hSyn-Cre is higher than 2 × 10^13^ genomic copies per ml. The titer of other viruses ranges from 2 × 10^12^ to 2 × 10^13^ genomic copies per ml.

### Stereotactic Surgery

Mice (8-20 weeks old) were anesthetized with 1%-2% isoflurane and mounted on a stereotaxic device (Kopf Instruments Model 1900). Viruses were delivered into brains through a glass capillary using nanoinjector (World Precision Instruments, Nanoliter 2000).

For investigating infanticide-induced c-Fos expression in MPOA-connected brain regions, 50 nl AAV1-hSyn-Cre (titer >2X10^13^) and 50 nl AAV2-hSyn-DIO-mCherry were mixed and injected into unilateral MPOA (AP: 0 mm, ML: -0.3 mm, DV: -4.95 mm) of SW Ai6 female mice.

For recording Ca^2+^ signal of MPOA^Esr1^ or BNSTpr^Esr1^ cells, 300 nl AAV2-CAG-Flex-GCaMP6f was injected into unilateral MPOA (AP: 0 mm, ML: -0.3 mm, DV: -4.95 mm) or BNSTpr (AP: -0.45 mm, ML: -0.9 mm, DV: -3.6 mm) of heterozygous virgin Esr1-2A-Cre female in SW background. For recording Ca^2+^ signals of BNSTpr^MPOA^ cells, 100 nl AAV1-hSyn-Cre (titer >2X10^13^) and 100 nl AAV2-hSyn-DIO-mCherry were mixed and injected into unilateral MPOA (AP: 0 mm, ML: -0.3 mm, DV: -4.95 mm), at the same time 300 nl AAV2-CAG-Flex-GCaMP6f was injected unilaterally into BNSTpr (AP: -0.45 mm, ML: - 0.9 mm, DV: -3.6 mm) of WT virgin female in SW background. After virus injection, a 400-µm optical fiber assembly (Thorlabs, FR400URT, CF440) was inserted 300 µm above the virus injection site and secured on the skull using adhesive dental cement (C&B Metabond, S380). Recording started at least 3 weeks after surgery.

For chemogenetically activating MPOA-connected cells in various brain regions, adult female mice were screened prior to surgery to ensure no spontaneous infanticide. Then, for each of the seven brain regions (BNSTpr, MeApd, PVT, PVN, VMHvl, PMv and SUM), 100 nl AAV1-hSyn-Cre (titer >2X10^13^) and 100 nl AAV2-hSyn-Flex-GFP were mixed and injected into bilateral MPOA (AP: 0 mm, ML: ±0.3 mm, DV: -4.95 mm), and at the same time AAV2-hSyn-DIO-hM3Dq-mCherry was injected into bilateral BNSTpr (AP: -0.45 mm, ML: ±0.9 mm, DV: -3.6 mm; 300 nl/side), MeApd (AP: -2.0 mm, ML: ±2.25 mm, DV: -4.6 mm; 200 nl/side), PVT (AP: -0.96 mm, ML: ±0.2 mm, DV: -3.17 mm; 100 nl/side), PVN (AP: -0.6 mm, ML: ±0.3 mm, DV: -4.3 mm; 100 nl/side), VMHvl (AP: -1.8 mm, ML: ±0.75 mm, DV: -5.6 mm; 50 nl/side), PMv (AP: -2.35 mm, ML: ±0.5 mm, DV: -5.6 mm; 200 nl/side) or SUM (AP: -3.06 mm, ML: ±0.4 mm, DV: -4.7 mm; 100 nl/side). For control females, AAV2-hSyn-DIO-mCherry instead of AAV2-hSyn-DIO-hM3Dq-mCherry was injected into the target region.

For chemogenetically inhibiting BNSTpr^Esr1^ neurons, adult heterozygous virgin Esr1-2A-Cre female in SW background were used for surgery. AAV2-hSyn-DIO-hM4Di-mCherry was injected bilaterally into BNSTpr (AP: -0.45 mm, ML: ±0.9 mm, DV: -3.6 mm; 300 nl/side). For control group, AAV2-hSyn-DIO-mCherry of the same volume was bilaterally injected into BNSTpr.

For optogenetically activating BNSTpr^MPOA^ neurons, 100 nl AAV1-hSyn-Cre (titer >2X10^13^) and 100 nl AAV2-hSyn-Flex-GFP were mixed and injected bilaterally into MPOA (AP: 0 mm, ML: ±0.3 mm, DV: -4.95 mm), and at the same time AAV2-EF1a-DIO-ChR2-EYFP was injected bilaterally into BNSTpr (AP: -0.45 mm, ML: ±0.9 mm, DV: -3.6 mm; 300 nl/side) of adult WT C57BL/6 females. For optogenetically activating BNSTpr^Esr1^ neurons, adult heterozygous virgin Esr1-2A-Cre female of C57BL/6 and SW background were used for surgery. SW adult female mice were screened prior to surgery to ensure no spontaneous infanticide. AAV2-EF1a-DIO-ChR2-EYFP was injected bilaterally into BNSTpr (AP: -0.45 mm, ML: ±0.9 mm, DV: -3.6 mm; 300 nl/side). During the surgery and after virus injection, two 200-µm optical fibers (Thorlabs, FT200EMT, CFLC230) were inserted 500 µm above the virus injection sites, one on each side, and secured on the skull using adhesive dental cement (C&B Metabond, S380).

For anterograde tracing of BNSTpr^Esr1^ and MPOA^Esr1^ neurons, 50 nl AAV2-hSyn-FLEX-GFP was injected unilaterally into BNSTpr (AP: -0.45 mm, ML: -0.9 mm, DV: -3.6 mm) or MPOA (AP: 0 mm, ML: -0.3 mm, DV: -4.95 mm) of heterozygous virgin Esr1-2A-Cre female in SW background.

For examining the synaptic connection from BNSTpr^Esr1^ cells to MPOA^Esr1^ cells, AAV2-EF1a-DIO-ChR2-EYFP was bilaterally injected into BNSTpr (AP: -0.45 mm, ML: ±0.9 mm, DV: -3.6 mm; 300 nl/side), and at the same time AAV2-hSyn-DIO-mCherry was injected bilaterally into MPOA (AP: 0 mm, ML: ±0.3 mm, DV: -4.95 mm; 300 nl/side). For MPOA^Esr1^ to BNSTpr^Esr1^ projection, AAV2-EF1a-DIO-ChR2-EYFP was bilaterally injected into MPOA (AP: 0 mm, ML: ±0.3 mm, DV: -4.95 mm; 300 nl/side), and at the same time AAV2-hSyn-DIO-mCherry was bilaterally injected into BNSTpr (AP: -0.45 mm, ML: ±0.9 mm, DV: -3.6 mm; 300 nl/side). All mice were heterozygous virgin Esr1-2A-Cre female in SW background.

For examining the intrinsic properties of MPOA^Esr1^ and BNSTpr^Esr1^ cells, AAV2-hSyn-FLEX-GFP virus was bilaterally injected into MPOA (AP: 0 mm, ML: ±0.3 mm, DV: -4.95 mm; 300 nl/side) and BNSTpr (AP: -0.45 mm, ML: ±0.9 mm, DV: -3.6 mm; 300 nl/side) in the same animal. All mice were heterozygous virgin Esr1-2A-Cre female in SW or C57BL/6 background.

### Fiber photometry

Fluorescence signals were acquired with a fiber photometry system as described previously ^26,51^. A 390-Hz sinusoidal blue LED light (30 mW; LED light: M470F1; LED driver: LEDD1B; from Thorlabs) were bandpass filtered (passing band: 472 ± 15 nm, FF02-472/30-25, Semrock) and delivered to the brain to excite GCaMP6f. The emission light traveled back through the same optic fiber, bandpass filtered (passing bands: 535 ± 25 nm, FF01-535/505, Semrock), passed through an adjustable zooming lens (Thorlab, SM1NR01 and Edmund optics, #62-561), detected by a Femtowatt Silicon Photoreceiver (Newport, 2151) and recorded using a real-time processor (RP2, TDT). The envelope of the 390-Hz signals reflected the intensity of the GCaMP and was extracted in real time using a custom TDT program. The signal was low pass filtered with a cut-off frequency of 10 Hz. The blue LED was adjusted at the tip of the optical fiber to 30 µW. The baseline fluorescence was set around 1 (arbitrary unit) for all animals by adjusting the zooming lens attached to the photoreceiver.

For fiber photometry recording of MPOA^Esr1^ neurons, 11 females were injected with AAV2-CAG-Flex-GCaMP6f, and 5 out of 11 females showed infanticide 3 weeks after surgery and were used for recording. During the first recording session, animals were left alone in their home cage for around 10 minutes, then a P1-P5 pup was introduce at a location distant from the nest. After naturally occurring infanticide, the pup was removed and euthanized. A total of 3-5 pups were introduced during the recording session with each for approximately 1-2 minutes. After the recording session with pups, a group housed adult Balb/c male mouse and then an adult Balb/c female mouse was introduced into the cage of the recording female mouse, each for 10 minutes with 10 minutes in between. After recording in hostile virgin state, females were exposed to pups more than half an hour each day for 1-2 weeks. During the first 3-7 days of pup sensitization, pups were presented under a cup to prevent infanticide. Once females stopped infanticide, they were allowed to freely interact with 3-5 pups for another 2-7 days until they quickly retrieved all pups back to nest on two consecutive days. Then the Ca^2+^ responses were recorded again. During the recording, 3-5 pups were introduced into the cage far from the nest for approximately 10 minutes. Then, adult male and female intruders were introduced sequentially in the same way as the recording under hostile virgin state. After completing both recording sessions, each female was paired with an adult male until becoming visibly pregnant, then male mice were removed. On postpartum day 2 or 3, mothers were recorded with the same procedure as that in maternal virgin state.

For fiber photometry recording of BNSTpr^Esr1^ neurons, 14 females that were injected with AAV2-CAG-Flex-GCaMP6f showed the proper virus expression and fiber placement. 8 out of 14 females showed infanticide and 7 out of 8 females became mothers. So, 7 females were recorded from virgin to lactating state. For fiber photometry recording of BNSTpr^MPOA^ neurons, 6 females were recorded from virgin to lactating state. The recording procedure was the same as that for MOPA^Esr1^ cells.

To analyze the recording data, the MATLAB function “msbackadj” with a moving window of 25% of the total recording duration was first applied to obtain the instantaneous baseline signal. The ΔF/F was then calculated as (F_raw_ − F_baseline_)/F_baseline_ subtracting the mean value during the period before intruder introduction. The PETHs were constructed by aligning the ΔF/F to the onset of each trial of a behavior, averaging across all trials for each animal and then averaging across animals. For each recording session, the responses during a behavior or no interaction period were calculated as the average ΔF/F during all trials of a behavior. Responses during the first contact was calculated as the maximum ΔF/F during the first contact period.

### Immunohistochemistry and imaging analysis

Mice were perfused with 1 X PBS followed with 4% PFA. Brains were dissected, post-fixed in 4% PFA overnight at 4°C, rinsed with 1 X PBS and dehydrated in 30% sucrose for 12-16 hours. 30 µm sections were cut on a Leica CM1950 cryostat. For brain wide c-Fos staining, every one in three brain slices of the whole brain were collected. For Esr1 staining, every one in three brain slices of BNSTpr region were collected. Then, free-floating brain slices were rinsed with PBS (3 × 10 min) and PBST (0.1% Triton X-100 in PBS, 1 × 30 minutes) at room temperature, followed with 1 hour blocking in 10% normal donkey serum at room temperature. First antibody (Guinea pig anti-Fos, 1:500 dilution, Synaptic Systems, Cat. # 226005; Rabbit anti-Esr1, 1:3000 dilution, Millipore, Cat. # 06-935, Lot # 3243424) was diluted in PBST with 3% normal donkey serum and incubated overnight (12-16 hours) at 4°C. Brain slices were then washed three times with PBST, each for 10 minutes. Then, brain slices were incubated with a secondary antibody (Secondary antibody for c-Fos staining, Cy3-Goat anti-Guinea pig, 1:500 dilution, Jackson ImmunoResearch, Cat. # 706-165-148; Secondary antibody for Esr1 staining, Cy5-Donkey anti-rabbit, 1:500 dilution, Jackson ImmunoResearch, Cat. # 711-175-152) for 2 hours at room temperature. Then brain slices were washed three times with PBST, each for 10 minutes. Lastly, brain slices were rinsed with 1 X PBS and mounted on superfrost slides (Fisher Scientific, 12-550-15), dried 10 minutes room temperature and cover slipped in 50% glycerol containing DAPI (Invitrogen, Cat. #00-4959-52). Images were acquired using a slide scanner (Olympus, VS120) or a confocal microscope (Zeiss LSM 700 microscope). Brain regions were identified based on Allen Mouse Brain Atlas, and cells were counted manually using ImageJ.

For the density of Fos^+^ cells (Fig. 1b), Fos^+^ cells in each brain region were counted, then divided by the area of each brain region to get the density of Fos^+^ cells in each brain region. For the Fos^+^Tracer^+^/Tracer^+^ cells in each brain region (Fig. 1c), the Fos^+^Tracer^+^ and Tracer^+^ cells in each brain region were counted, then use Fos^+^Tracer^+^ cell number divided by Tracer^+^ cells numbers to get the activated fraction. For relative projection density (Extended Data Figs. 7e and 7j), average fluorescence intensity in each region containing presynaptic GFP+ punctates was calculated, then normalized by the average fluorescence intensity of the start region (Extended Data Figs. 7e normalized by BSNTpr, 7j normalized by MPOA).

### Pharmacogenetic activation and inactivation

For pharmacogenetic activation of MPOA-connected cells in BNSTpr, MeApd, PVT, PVN, VMHvl, PMv and SUM, 8 females were tested for each brain region. On the day of behavior test, sterile saline or CNO (1 mg/kg) was injected intraperitoneally 30 min prior to behavioral assays. During the 10 minutes behavior test, 3-4 P1-P5 pups were scattered in the testing female home cage. After the test, wounded pups were removed and euthanized.

For pharmacogenetic inhibition of BNSTpr^Esr1^ neurons assay, 13 hM4Di and 13 mCherry control females were tested in the virgin state. 11 out of 13 hM4Di females successfully became mothers and all showed maternal aggression towards adult male and female intruders. 9 out of 13 mCherry control females successfully became mothers and 8 showed maternal aggression. All animals were first tested after saline injection and then after CNO (1 mg/kg) injection as females often stopped spontaneous infanticide after showing maternal behaviors induced by CNO injection. For behavior test in virgin females, 30 min after saline or CNO injection, 3-4 P1-P5 pups were scattered in the testing female’s home cage for 10 minutes. For tests in lactating females (postpartum day 3 and day 4), 30 min after saline or CNO injection, 5 pups were introduced into the home cage far from the nest and recorded 10 minutes. After the pup section, an adult group housed Balb/c male and then a Balb/c female was introduced into the cage for 10 minutes with 10 minutes in between. Animal behaviors were recorded from both top and side using two synchronized cameras (Edmund, stock #89533) controlled by StreamPix (NORPIX) at 25 frames/sec.

### Optogenetic neural activation

For BNSTpr^MPOA^ optogenetic activation, 11 WT C57BL/6 virgin adult female mice were injected with ChR2 virus, 8 WT C57BL/6 virgin adult female mice were injected with GFP virus. For BNSTpr^Esr1^ optogenetic activation, 8 Esr1-2A-Cre C57BL/6 and 5 Esr1-2A-Cre SW virgin adult female mice were injected with ChR2 virus, and 8 Esr1-2A-Cre C57BL/6 virgin adult female mice were injected with GFP virus. Esr1-2A-Cre SW virgin adult female mice were pre-screened and only females that did not show infanticide were used for surgery.

Three weeks after surgery, the implanted optic fiber assembly was coupled to a patch cord using a zirconia split sleeve to deliver 473 nm laser pulses to the brain. The laser pulses were controlled by TTL signals generated using a RP2 processor (TDT). For each test session, 9 trials of 20 ms, 20 Hz laser pulses were delivered, each lasting for 20 seconds with 40 seconds in between. The first pulse train started when the testing female investigated the intruder. Each test session started with sham (laser off) stimulation followed by stimulations with light intensities at 0.5 mW, 1 mW, 2 mW, 3 mW, 4 mW and 5 mW. During pup session, 3-4 P1-P5 pups were scatted around the testing female home cage and the light was delivered according to the protocol described above. Once the females attacked a pup, the wounded pup was removed and euthanized within 1 minute. For adult female and male sessions, a group housed adult Balb/c female or male was introduced and the light trials were delivered based on the stimulation protocol. For BNSTpr^Esr1^ group, after completing the test in virgin state, each female was paired with an adult male until becoming visibly pregnant and tested again on postpartum day 3 or 4. Ten minutes before the test session, all pups were removed. During the test, the pups were returned to the female’s cage at a location far from the nest. Then, the light trials were delivered as described above.

For RTPP test, sham or 3 mW light pulses were delivered whenever the animal entered the pre-designated stimulation chamber and terminated when the animal moved out of the chamber. Each test lasted for 10 minutes. For EPM test, mice were habituated to the test area for two days, 20 minutes a day. During the test, sham light was delivered for the first 20 minutes followed by 3 mW light pulses for 20 minutes. The body center of the animal was tracked with DLC and used for calculating the time animal spent in each chamber in RTPP test and the open/closed arms in EPM test.

### Behavioral analyses

Behaviors were analyzed frame-by-frame using a custom software written in MATLAB (https://pdollar.github.io/toolbox/). Approaching pup is when the testing female faces and walks straight to the pup. Investigating pup is when the female’s nose closely contacts any body parts of a pup. Attacking pup is defined as biting of a pup and confirmed by the wounds. Retrieving pup is defined as the moment the female lift the pup using jaw to the moment when the pup is dropped in or around the nest. Grooming pup is defined as close interaction between the female and pup that is accompanied by rhythmic up and down head movement of the female and displacement of the pup. Investigating female or male is defined as nose-to-face, nose-to-trunk, or nose-to-urogenital contact. Social grooming is defined as licking or grooming the head or neck area of the adult intruder. Mounting is when the testing female mouse clasps onto the flank of the adult intruder, establishes an on-top position and moves its pelvis rhythmically. Attacking female or male is defined as lunging, biting and fast movements connecting these behaviors.

### *In vitro* electrophysiological recordings

For *in vitro* whole cell patch-clamp recording, mice were anesthetized with isoflurane, and brains were removed and submerged in ice-cold cutting solution containing (in mM): 110 choline chloride, 25 NaHCO_3_, 2.5 KCl, 7 MgCl_2_, 0.5 CaCl_2_, 1.25 NaH_2_PO_4_, 25 glucose, 11.6 ascorbic acid and 3.1 pyruvic acid. 275-µm coronal sections were cut on a Leica VT1200s vibratome and incubated in artificial cerebral spinal fluid (ACSF) containing (in mM): 125 NaCl, 2.5 KCl, 1.25 NaH_2_PO_4_, 25 NaHCO_3_, 1 MgCl_2_, 2 CaCl_2_ and 11 glucoses at 34° C for 30 min and then at room temperature until use. The intracellular solution for current clamp recording contained (in mM): 145 K-gluconate, 2 MgCl_2_, 2 Na_2_ATP, 10 HEPES, 0.2 EGTA (286 mOsm, pH 7.2). The intracellular solution for the voltage clamp recording contained (in mM): 135 CsMeSO_3_, 10 HEPES, 1 EGTA, 3.3 QX-314 (chloride salt), 4 Mg-ATP, 0.3 Na-GTP and 8 sodium phosphocreatine (pH 7.3 adjusted with CsOH). The signals were acquired using MultiClamp 700B amplifier and digitized at 20 kHz with DigiData1550B (Molecular Devices, USA). The recorded electrophysiological data were analyzed using Clampfit (Molecular Devices) and MATLAB (Mathworks).

To determine the intrinsic excitability of MPOA^Esr1^ and BNSTpr^Esr1^ cells, we performed current clamp recording and injected 30 current steps ranging from -20 pA to 270 pA in 10 pA increments into the recorded cell. The total number of spikes during each 500-ms long current step was then used to construct F-I curve.

Voltage clamp recording was conducted for BNSTpr^Esr1^ and MPOA^Esr1^ neurons, and the membrane voltage was held at -70 mV for oEPSC recording, and at 0 mV for oIPSC recording. To activate ChR2-expressing axons, brief pulses of full field illumination (0.5 ms duration, 10 repeats) were delivered onto the recorded cell with a blue LED light (pE-300 white; CoolLED) at 10 s interval. TTX (1 µM), 4-AP (100 mM), Gabazine (2 µM) and strychnine (5 µM) were then applied to the bath solution to block GABAergic transmission and determine if the optogenetically evoked responses were monosynaptic. All drugs were perfused for at least 5 minutes prior to data acquisition.

### Statistics

All statistical analysis were performed using MATLAB and Prism software. Parametric tests, including paired *t test*, and unpaired *t test* were used if distributions passed Kolmogorov Smirnov tests for normality, a nonparametric Wilcoxon test was used if distributions not normally distributed. Fisher’s exact test was used when compare two nominal variables. For more than two groups repeated tests, statistical significances were determined using One-way ANOVA followed with Tukey correction. For comparing among multiple groups and multiple treatment conditions, Two-way ANOVA were used, and if comparing every row (or column) mean with every other row (or column) mean, Tukey correction was used. If comparing every row (or column) mean with the control row (or column) mean, Bonferroni correction was used. For all statistical tests, significance was measured against an alpha value of 0.05. For detailed statistical results, see statistic summary table.

### Data Availability and Code Availability

Source data are provided with this paper. Behavioral analysis code can be found (https://pdollar.github.io/toolbox/). Additional data and code relating to the paper are available from the corresponding author on reasonable request.

