## Supplementary Figures for "Antagonistic Neural Circuits Drive Opposing Behaviors towards the Young in Females"

#### Extended Data Figures and Legends

### Extended Data Figure 1

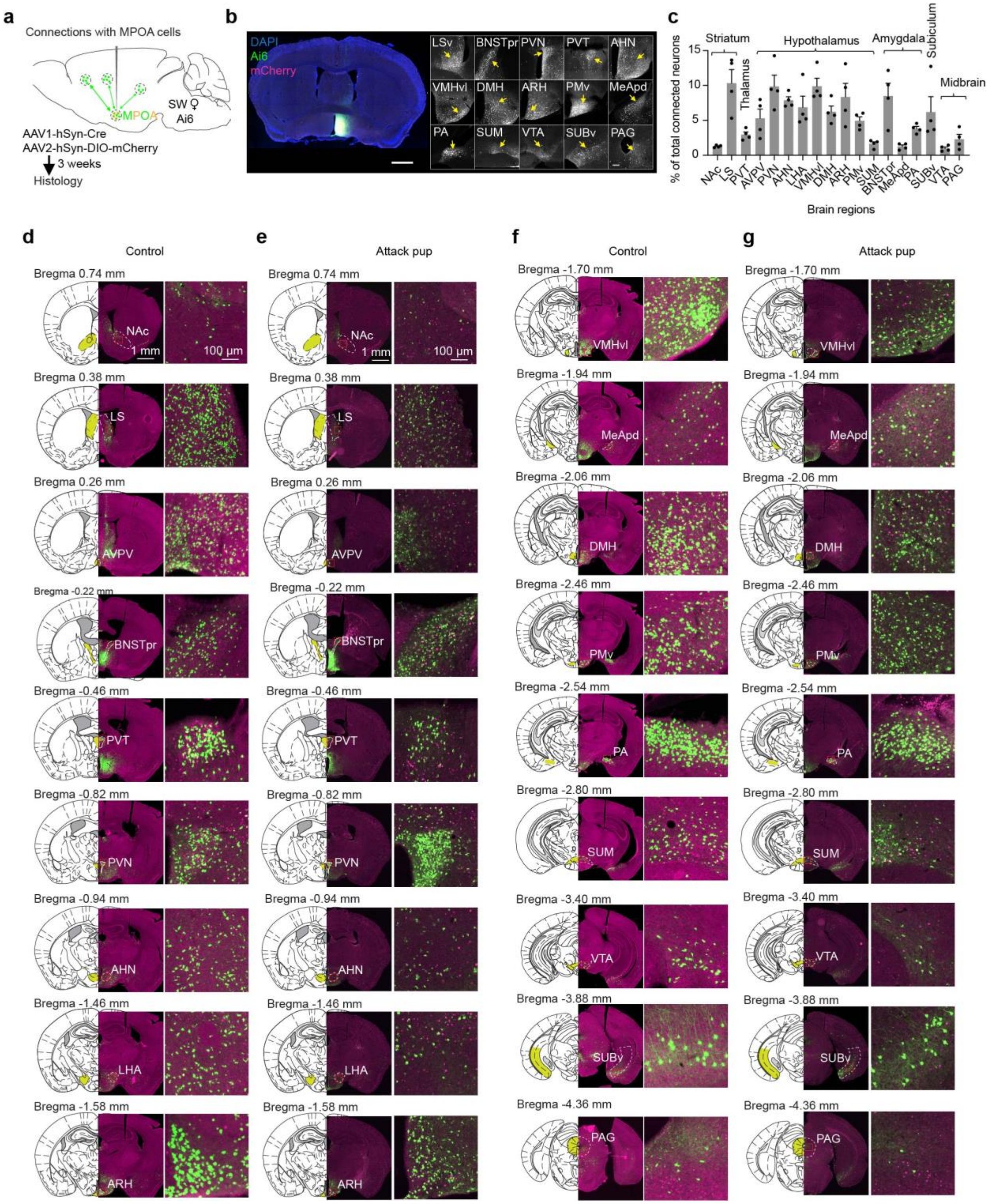

**Extended Data Fig. 1: Infanticide induces c-Fos in MPOA-connected cells in multiple brain regions**

**(a)** Experimental design to trace MPOA-connected cells throughout the brain using Ai6 female mice and high titer (> $1\times10^{13}$ vg/mL) AAV1-hSyn-Cre.

**(b)** Images from a representative animal showing the primary injection site in the MPOA and MPOA-connected cells in various brain areas. Scale bars, 1 mm (left) and 200 µm (right).

**(c)** Distribution of MPOA-connected neurons in various brain regions. All regions containing over 1% of total labeled cells are shown. n = 4. Error bars: SEM.

**(d-g)** Representative images in each of 18 regions showing c-Fos staining (magenta) and zsGreen expression in Ai6 female mice injected with AAV1-hSyn-Cre into MPOA. The females either showed infanticide (**e and g**) or alone in home cage (**d and f**).

Source data provided. Details of the statistical analyses and sample sizes can be found in Statistic Summary Table.

### Extended Data Figure 2

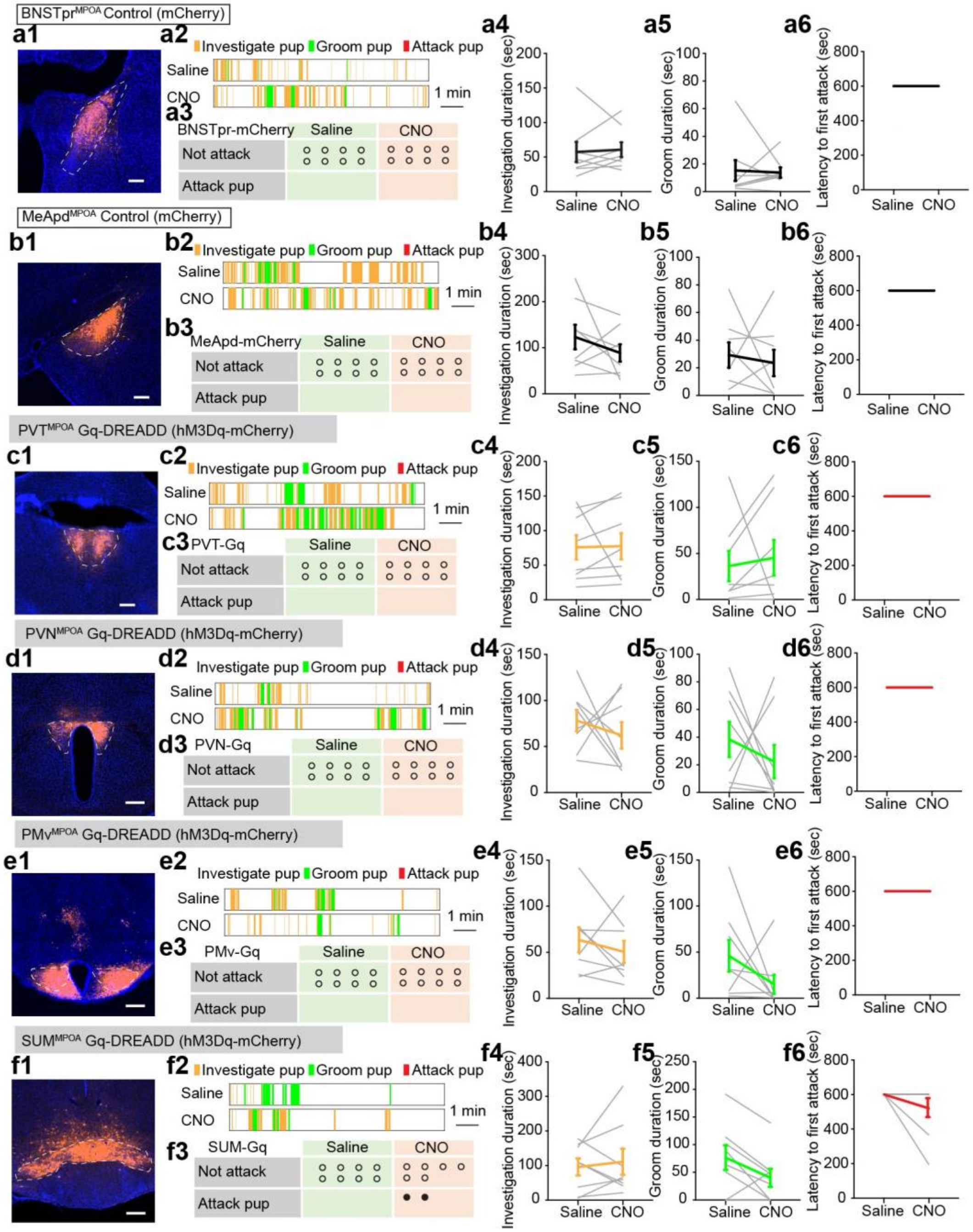


**Extended Data Fig. 2: Pup-directed behaviors after chemogenetic activation of MPOA-connected cells in PVT, PVN, PMv or SUM and mCherry controls for BNSTpr, MeApd regions in female mice.**

**(a1, b1, c1, d1, e1, f1)** Representative histology images showing hM3Dq-mCherry or mCherry (red) expression in the BNSTpr (**a1**, mCherry), MeApd (**b1**, mCherry), PVT (**c1**, hM3Dq-mCherry), PVN (**d1**, hM3Dq-mCherry), PMv (**e1**, hM3Dq-mCherry) and SUM (**f1**, hM3Dq-mCherry) after injecting AAV1-Syn-Cre in MPOA and Cre-dependent hM3Dq-mCherry or mCherry in each of the regions. Scale bars: 200 µm.

**(a2, b2, c2, d2, e2, f2)** Representative raster plots showing pup-directed behaviors after saline or CNO injection into animals expressing mCherry in MPOA-connected cells in BNSTpr (**a2**) or MeApd (**b2**), or animals expressing hM3Dq-mCherry in MPOA-connected cells in PVT (**c2**), PVN (**d2**), PMv (**e2**) or SUM (**f2**). Each raster lasts 10 min.

**(a3, b3, c3, d3, e3, f3)** Pup-directed aggressive behaviors after saline or CNO injection in female mice that express mCherry or hM3Dq-mCherry in MPOA-connected cells in BNSTpr (**a3**, mCherry), MeApd (**b3**, mCherry), PVT (**c3**, hM3Dq-mCherry), PVN (**d3**, hM3Dq-mCherry), PMv (**e3**, hM3Dq-mCherry) or SUM (**f3**, hM3Dq-mCherry). Each dot represents one mouse. n = 8 for each group.

**(a4, b4, c4, d4, e4, f4)** Duration of pup investigation did not differ between post-saline and post-CNO injections in any group as in **a1-f1**. Each gray line represents one animal. Colored line represents the group average. Error bars: ± SEM. *t-test*. n = 8 for each group.

**(a5, b5, c5, d5, e5, f5)** Duration of pup grooming did not differ between post-saline and post-CNO injections in any group as in **a1-f1.** Figure conventions as in **a4-f4**.

**(a6, b6, c6, d6, e6, f6)** Latency to first pup attack did not differ between post-saline and post-CNO injections in any group as in **a1-f1.** Figure conventions as in **a4-f4**.

Source data provided. Details of the statistical analyses and sample sizes can be found in Statistic Summary Table.

### Extended Data Figure 3

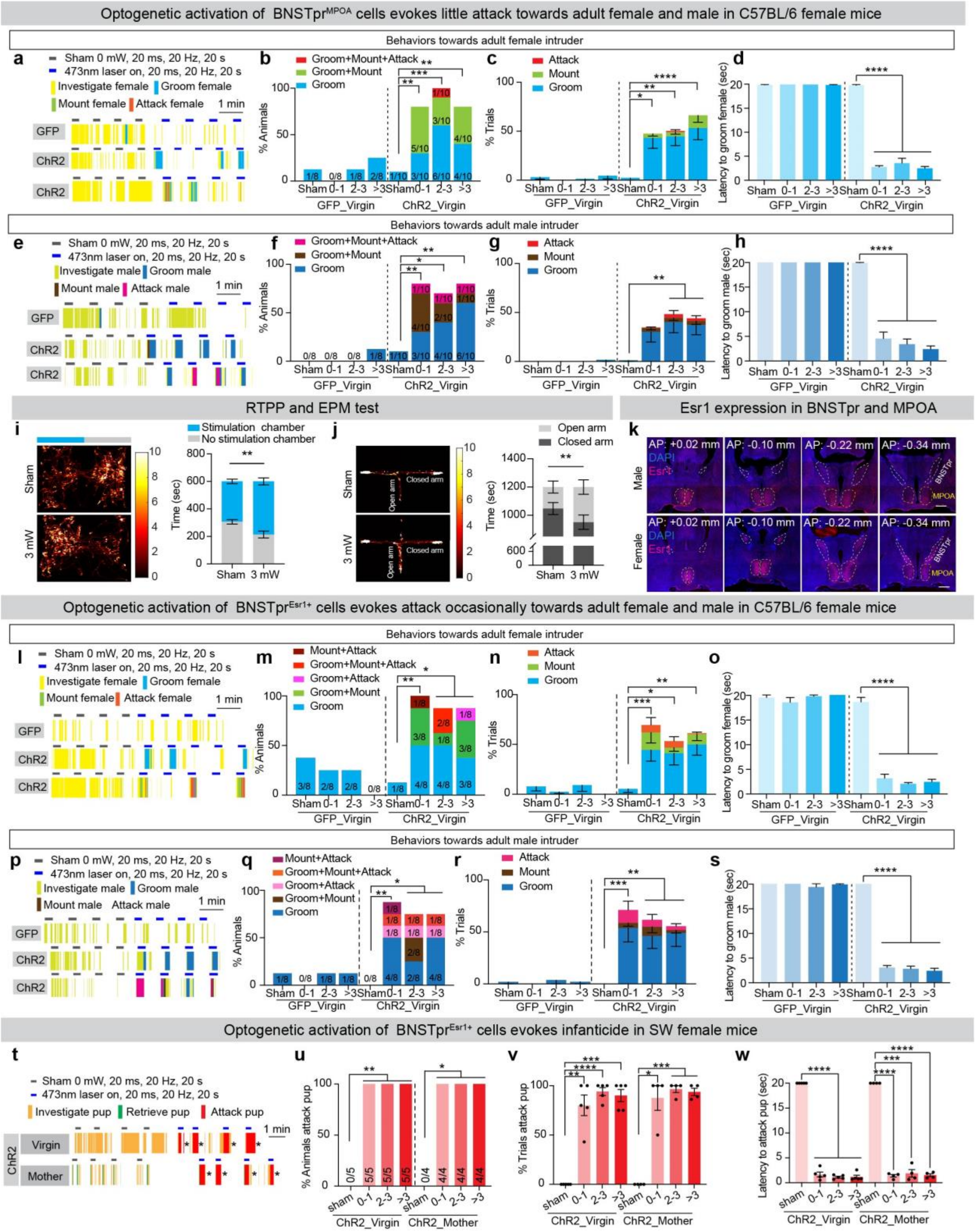

**Extended Data Fig. 3: Optogenetic activation of BNSTpr neurons elicits social grooming in female mice**

**(a)** Representative raster plots showing behaviors toward a female intruder during sham or laser stimulation of a $BNSTpr^{MPOA}$-GFP control mouse (top) and two $BNSTpr^{MPOA}$-ChR2 test mice (middle and bottom). Note that during light stimulation one ChR2 mouse showed only grooming while the other showed both grooming, brief male-style mounting and attacking.

**(b)** Percentage of animals showing each behavior towards the female intruder during $BNSTpr^{MPOA}$ manipulation. Fisher's exact test. **p < 0.01. ***p < 0.001.

**(c)** Percentage of $BNSTpr^{MPOA}$ sham or stimulation trials that animals show each type of behaviors towards a female intruder. Two-way ANOVA. *p < 0.05, **p < 0.01, ***p < 0.001. Error bars: SEM.

**(d)** The latency to groom the female intruder during $BNSTpr^{MPOA}$ stimulation. Two-way ANOVA. ****p < 0.0001. Error bars: SEM.

**(e-h)** Behavior changes towards an adult male intruder induced by optogenetic activation of $BNSTpr^{MPOA}$ cells in C57BL/6 WT female mice. Figure conventions as in **a-d**.

**(i)** Representative tracking results during the RTPP test (left) and the time spent in each chamber (right) with sham or 3 mW laser stimulation. n = 8 mice. Two-way ANOVA. **p < 0.01. Error bars: ± SEM.

**(j)** Representative tracking results during an EPM test (left) and the time spent in open and closed arms (right) with sham or 3 mW laser stimulation. n = 8 mice. Two-way ANOVA. ***p < 0.001. Error bars: ± SEM.

**(k)** Representative coronal sections showing Esr1 immunostaining in MPOA and BNSTpr in a male (top) and a female (bottom) mouse. Scale bars: 500 µm.

**(l-o)** Behavior changes towards an adult female intruder induced by optogenetic activation of $BNSTpr^{Esr1}$ cells in Esr1-2A-Cre C57BL/6 female mice. Figure conventions as those in **a-d**.

**(p-s)** Behavior changes towards an adult male intruder induced by optogenetic activation of $BNSTpr^{Esr1}$ cells in Esr1-2A-Cre C57BL/6 female mice. Figure conventions as those in **a-d**.

**(t)** Representative raster plots showing pup-directed behaviors during sham or light stimulation of $BNSTpr^{Esr1}$ cells in SW virgin and lactating female mice. *Remove the hurt pup and introduce a new pup.

**(u)** Percentage of SW $BNSTpr^{Esr1}$-ChR2 animals that attacked pups during sham and light stimulation. Fisher's exact test. *$p < 0.05$, **$p < 0.01$.

**(v)** Percentage of trials that the test animal attacked a pup. Each dot represents one mouse. Two-way. *$p < 0.05$, **$p < 0.01$, ***$p < 0.001$, ****$p < 0.0001$. Error bars: ± SEM.

**(w)** Average latency to attack after encountering the pup following laser stimulation. Two-way. ***$p < 0.001$, ****$p < 0.0001$. Error bars: ± SEM. Each trial lasts 20 s. If no attack occurred during a trial, 20 s was used as the latency value.

Source data provided. Details of the statistical analyses and sample sizes can be found in Statistic Summary Table.

### Extended Data Figure 4

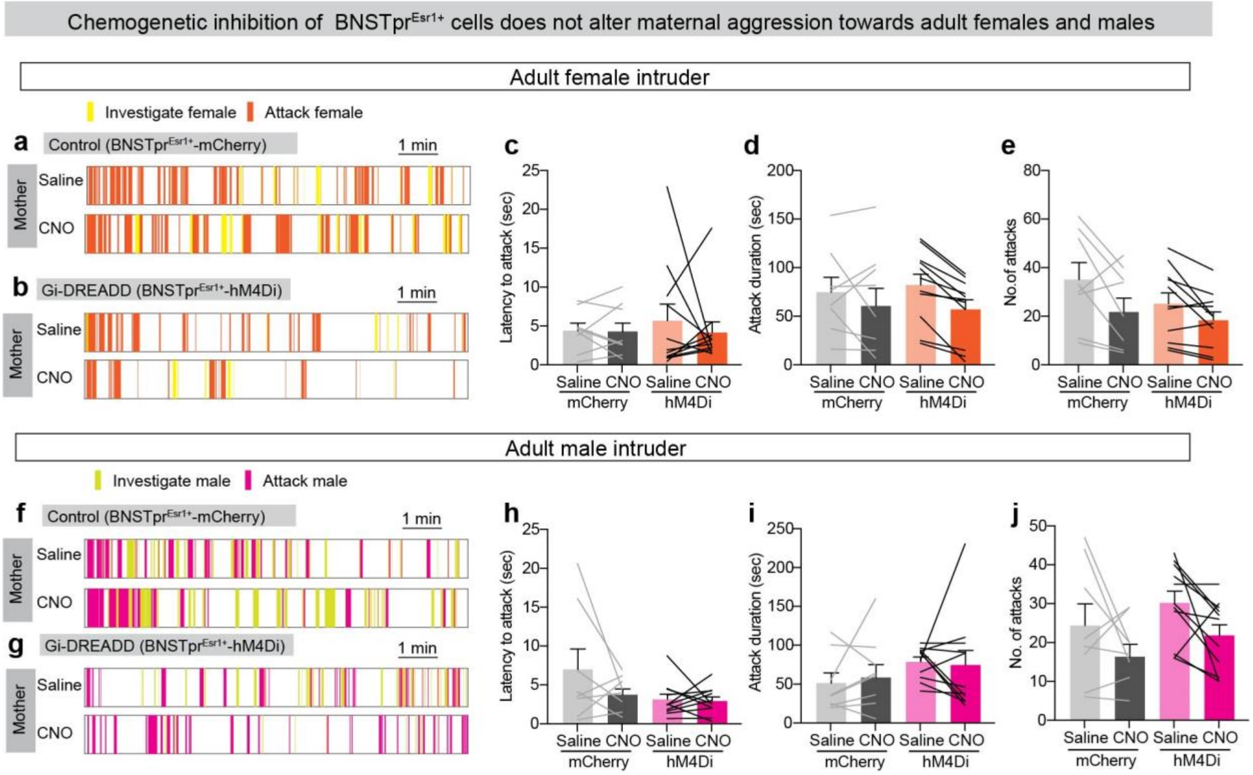


**Extended Data Fig. 4: BNSTpr$^{Esr1}$ neurons are not required for maternal aggression**

**(a and b)** Representative raster plots showing behaviors towards a female intruder of a lactating mCherry control female (**a**) and a lactating hM4Di test female (**b**) after saline or CNO injection.

**(c-e)** Latency to attack (**c**), attack duration (**d**) and number of attacks (**e**) towards a female intruder in lactating mCherry females and hM4Di females after saline or CNO injection. n= 8 mice for mCherry group and 11 for hM4Di group. Error bars: SEM.

**(f-j)** No behavioral change towards adult male intruders after chemogenetic inhibition of BNSTpr$^{Esr1}$ cells in Esr1-2A-Cre female mice. n= 8 mice for mCherry group and 11 for hM4Di group. Figure conventions as **a-e**.

Source data provided. Details of the statistical analyses and sample sizes can be found in Statistic Summary Table.

### Extended Data Figure 5

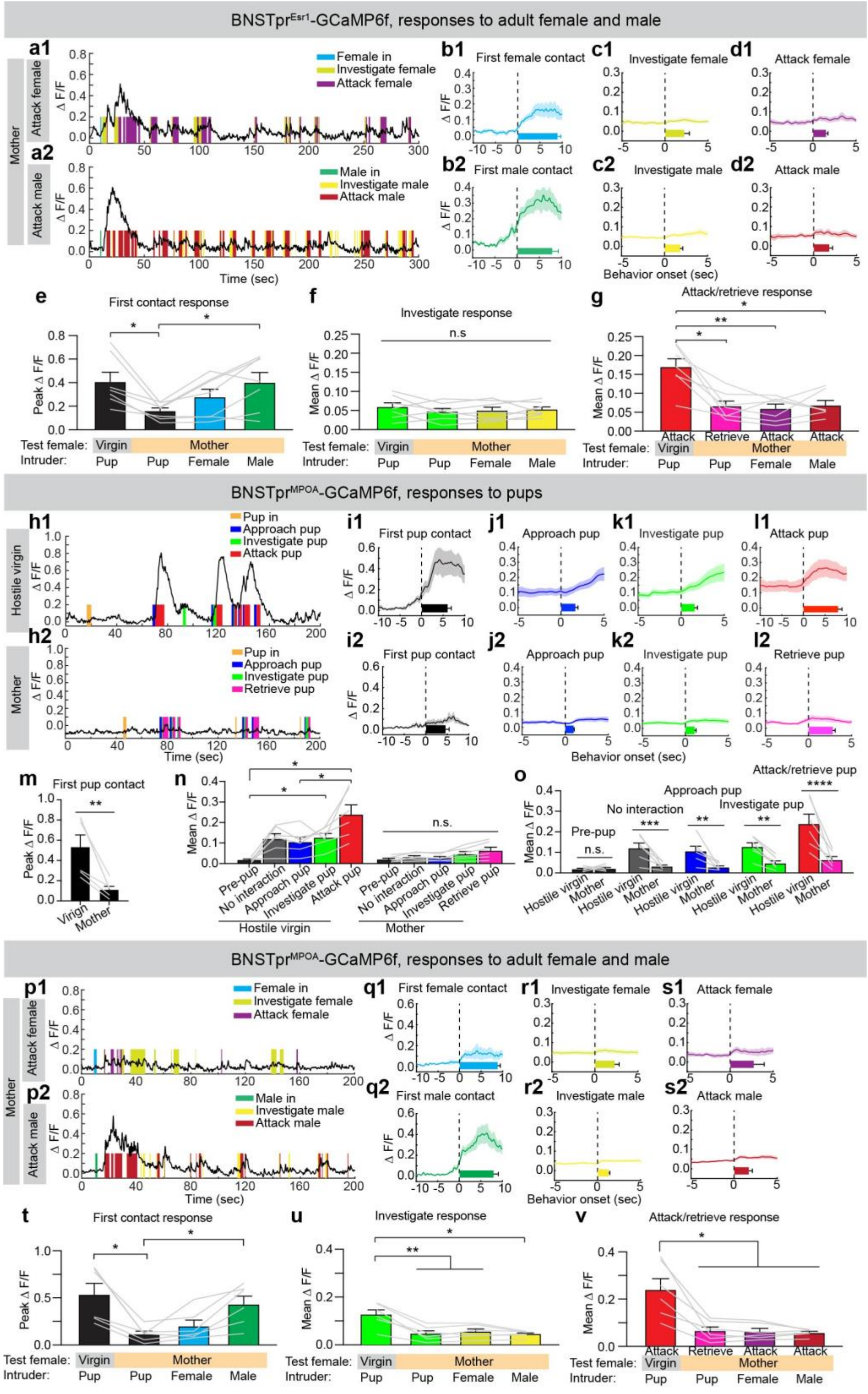

**Extended Data Fig. 5: $Ca^{2+}$ responses of $BNSTpr^{Esr1}$ cells to adult conspecifics, and $BNSTpr^{MPOA}$ cells response to various social stimuli.**

**(a)** Representative GCaMP6f recording (ΔF/F) traces of $BNSTpr^{Esr1}$ cells in a lactating SW female when interacting with an adult female intruder (**a1**) or an adult male intruder (**a2**). Color shades indicate various behaviors.

**(b)** PETHs of GCaMP6f signal (ΔF/F) of $BNSTpr^{Esr1}$ cells aligned to the onset of first contact with a female intruder (**b1**) or a male intruder (**b2**). n = 7 mice. Horizontal bars in the graph indicate the average duration of a behavior. Shades and error bars: SEM.

**(c and d)** PETHs of GCaMP6f signal (ΔF/F) of $BNSTpr^{Esr1}$ cells aligned to the onset of investigating or attacking an adult female (**c1 and d1**) or male (**c2 and d2**) intruder. n = 7 mice. Horizontal bars in the graph indicate the average duration of a behavior. Shades and error bars: SEM.

**(e-g)** Peak responses during the first contact (**e**), mean investigation responses (**f**) and mean attack or retrieve responses (**g**) of $BNSTpr^{MPOA}$ cells towards various intruders. Each gray line represents data from one animal. One-way ANOVA followed by Tukey's multiple comparison test, *$p < 0.05$, **$p < 0.01$, ***$p < 0.001$. Error bars: SEM.

**(h)** Representative GCaMP6f recording (ΔF/F) traces of $BNSTpr^{MPOA}$ cells during pup interaction from a hostile SW virgin female (**h1**) and a mother (**h2**). Color shades indicate various behaviors.

**(i-l)** PETHs of GCaMP6f signal (ΔF/F) of $BNSTpr^{MPOA}$ cells aligned to the onset of various behaviors in hostile virgin females **(i1-l1**), and mothers (**i2-l2**). n = 6 mice. Horizontal bars indicate the average duration of behaviors. Shades and error bars: SEM.

**(m)** Peak responses (ΔF/F) of $BNSTpr^{MPOA}$ cells during the first pup contact. Paired *t-test*. **$p < 0.01$, Error bars: SEM.

**(n-o)** Mean GCaMP6f signal (ΔF/F) of $BNSTpr^{MPOA}$ cells during pre-pup period and various pup-directed behaviors in hostile virgin females and mothers. Two-way ANOVA with Bonferroni's multiple comparisons test. **$p < 0.01$, ***$p < 0.001$ ****$p < 0.0001$, Error bars: SEM.

**(p)** GCaMP6f recording (ΔF/F) traces of $BNSTpr^{MPOA}$ cells of a SW mother during interactions with an adult female intruder (**p1**) and an adult male intruder (**p2**). Color shades indicate various behaviors.

**(q-s)** PETHs of GCaMP6f signal ($\Delta F/F$) of BNSTpr$^{MPOA}$ cells aligned to the onset of first contact with an adult female intruder (**q1**) or an adult male intruder (**q2**)**,** adult female investigation (**r1**)**,** adult male investigation (**r2**)**,** attack female (**s1**) and attack male (**s2**). Horizontal bars indicate the average duration of behaviors. Shades and error bars: SEM.
**(t-v)** Bar graphs showing the peak response during first contact (**t**), mean responses during investigation (**u**) and mean responses during attack or retrieve (**v**) of BNSTpr$^{MPOA}$ cells towards various social stimuli. One-way ANOVA followed by Tukey's multiple comparison test, $^{*}p < 0.05$, $^{**}p < 0.01$. Error bars: SEM.
Source data provided. Details of the statistical analyses and sample sizes can be found in Statistic Summary Table.

### Extended Data Figure 6

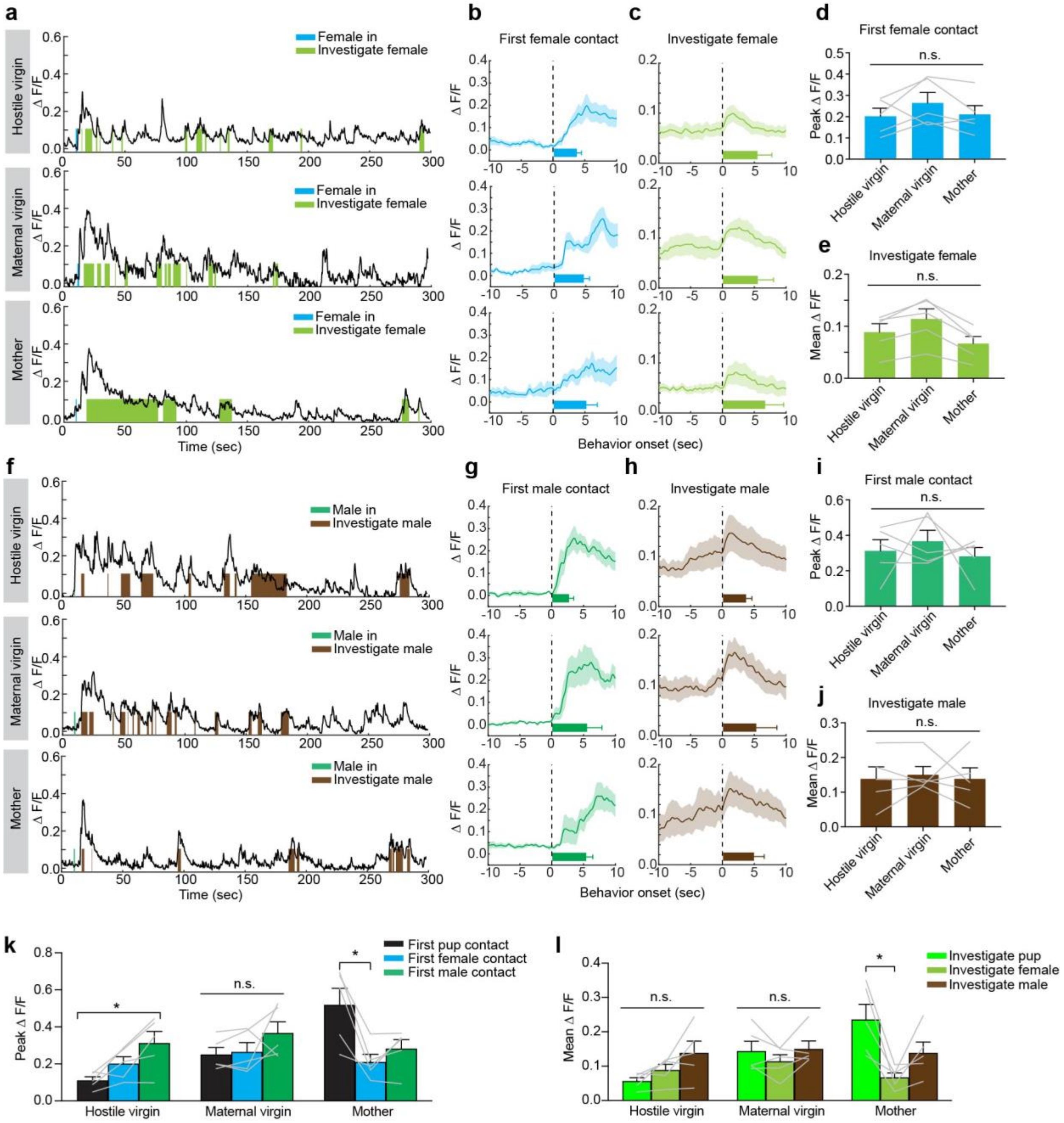


**Extended Data Fig. 6: Comparison of MPOA$^{Esr1}$ cells response to adult male and adult female mice.**

**(a)** Representative GCaMP6f recording (ΔF/F) traces of MPOA$^{Esr1}$ cells from a hostile virgin female (top), a maternal virgin female (middle) and a mother (down) when interacting with an adult female intruder. Color shades indicate various behaviors.

**(b and c)** PETHs of GCaMP6f signal (ΔF/F) aligned to the first contact with a female intruder (**b**) and the onset of investigation (**c**). n = 5 mice. Shading: SEM. Horizontal bars show the duration of each behavior. Error bars: SEM.

**(d and e)** Mean peak responses of MPOA$^{Esr1}$ cells during the first contact with a female intruder (**d**) and the mean response during female investigation (**e**). Error bars: SEM.

**(f)** Representative GCaMP6f recording (ΔF/F) traces from a hostile virgin female (top), a maternal virgin female (middle) and a mother (bottom) when interacting with an adult male intruder. Color shades indicate various behaviors.

**(g and h)** PETHs of GCaMP6f signal (ΔF/F) aligned to the first contact with a male intruder (**g**) and the onset of investigation (**h**). n = 5 mice. Shading: SEM. Horizontal bars show the duration of each behavior. Error bars: SEM.

**(i and j)** Mean peak responses during the first contact with a male intruder (**i**) and the mean response during male investigation (**j**). Error bars: SEM.

**(k)** The peak responses during the first contact with pups, females or males in females under different reproductive states. Two-way ANOVA. *$p < 0.05$. Error bars: SEM.

**(l)** The mean responses of MPOA$^{Esr1}$ cells during investigating pups, females and males in female mice under different reproductive states. Two-way ANOVA. *$p < 0.05$. Error bars: SEM.

Source data provided. Details of the statistical analyses and sample sizes can be found in Statistic Summary Table.

### Extended Data Figure 7

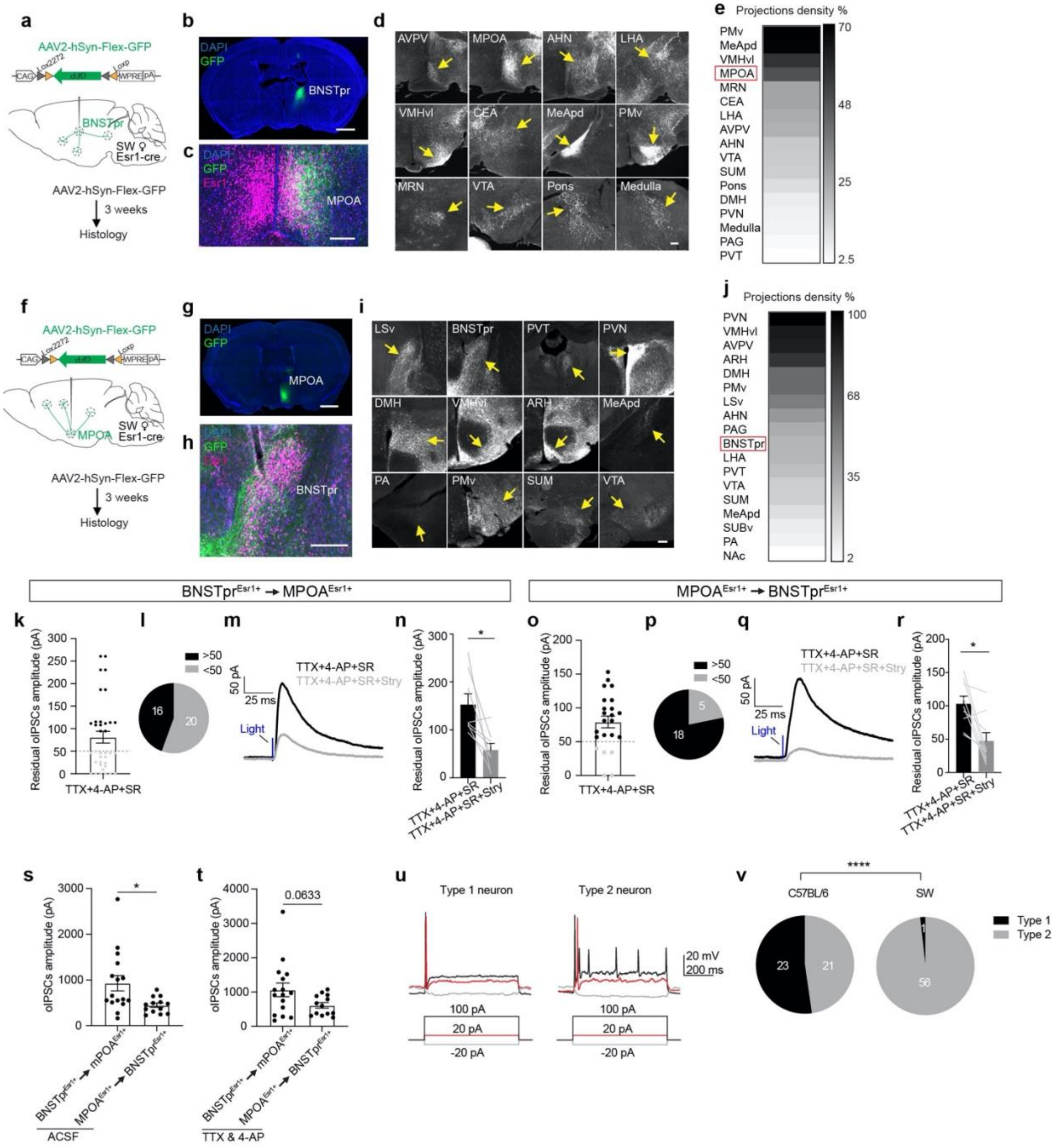


**Extended Data Fig. 7: The brain-wide projection pattern of BNSTpr$^{Esr1}$ and MPOA$^{Esr1}$ cells, and additional slice recording data from BNSTpr$^{Esr1}$ cells and MPOA$^{Esr1}$ cells.**

**(a)** Experimental design to examine the projection pattern of BNSTpr$^{Esr1}$ cells.

**(b)** An image showing the GFP (green) expression in BNSTpr$^{Esr1}$ cells. Blue, DAPI. Scale bar: 1mm.

**(c)** An image showing GFP (green) fiber terminals in the MPOA from BNSTpr$^{Esr1}$ cells and Esr1 staining (magenta). Blue, DAPI. Scale bar: 200 µm.

**(d)** Representative images showing regions containing dense GFP fibers from BNSTpr$^{Esr1}$ cells. Scale bar: 200 µm.

**(e)** Heatmap showing the brain wide projection density of BNSTpr$^{Esr1}$ cells. The value in each region is normalized by the fluorescence intensity in the BNSTpr. n = 4 mice.

**(f-j)** Brain-wide projection pattern of MPOA$^{Esr1}$ cells. n = 4 mice (**j**). Figure conventions as **a-e**.

**(k and o)** Amplitude of oIPSCs of MPOA$^{Esr1}$ cells (**k**) and BNSTpr$^{Esr1}$ cells (**o**) after bath application of TTX, 4-AP and gabazine mixture. n = 36 MPOA$^{Esr1}$ cells from 8 mice (**k**) and 23 BNSTpr$^{Esr1}$ cells from 9 mice (**o**). Error bars: SEM.

**(l and p)** Pie charts showing MPOA$^{Esr1}$ cells (**l**) or BNSTpr$^{Esr1}$ cells (**p**) that have a residual oIPSC amplitude smaller or larger than 50 pA after bath application of TTX + 4-AP + gabazine.

**(m and q)** Representative recording traces showing oIPSCs of MPOA$^{Esr1}$ cells (**m**) and BNSTpr$^{Esr1}$ cells (**q**) before and after strychnine application.

**(n and r)** Amplitude of oIPSCs of MPOA$^{Esr1}$ (**n**) and BNSTpr$^{Esr1}$ cells (**r**) before and after applying strychnine. Paired t test, *p<0.05. n = 8 MPOA$^{Esr1}$ cells from 4 mice (**n**) and 9 BNSTpr$^{Esr1}$ cells from 4 mice (**r**). Error bars: SEM.

**(s and t)** Comparison of the oIPSC amplitude of MPOA$^{Esr1}$ cells and BNSTpr$^{Esr1}$ cells in ACSF (**s**) or after application of TTX and 4-AP (**t**). Unpaired t test, *p<0.05. n= 16 MPOA$^{Esr1}$ cells from 4 mice (**s**) and 13 BNSTpr$^{Esr1}$cells from 5 mice (**t**). Error bars: ± SEM.

**(u)** Representative traces showing spiking patterns of type I (left) and type II (right) BNSTpr$^{Esr1}$ cells with current injections.

**(v)** Pie charts showing the percentage of type I and type II BNSTpr$^{Esr1}$ neurons in C57BL/6 (left) and SW (right) virgin female mice. Fisher's exact test, ****p<0.0001.

Source data provided. Details of the statistical analyses and sample sizes can be found in Statistic Summary Table.

### Extended Data Figure 8

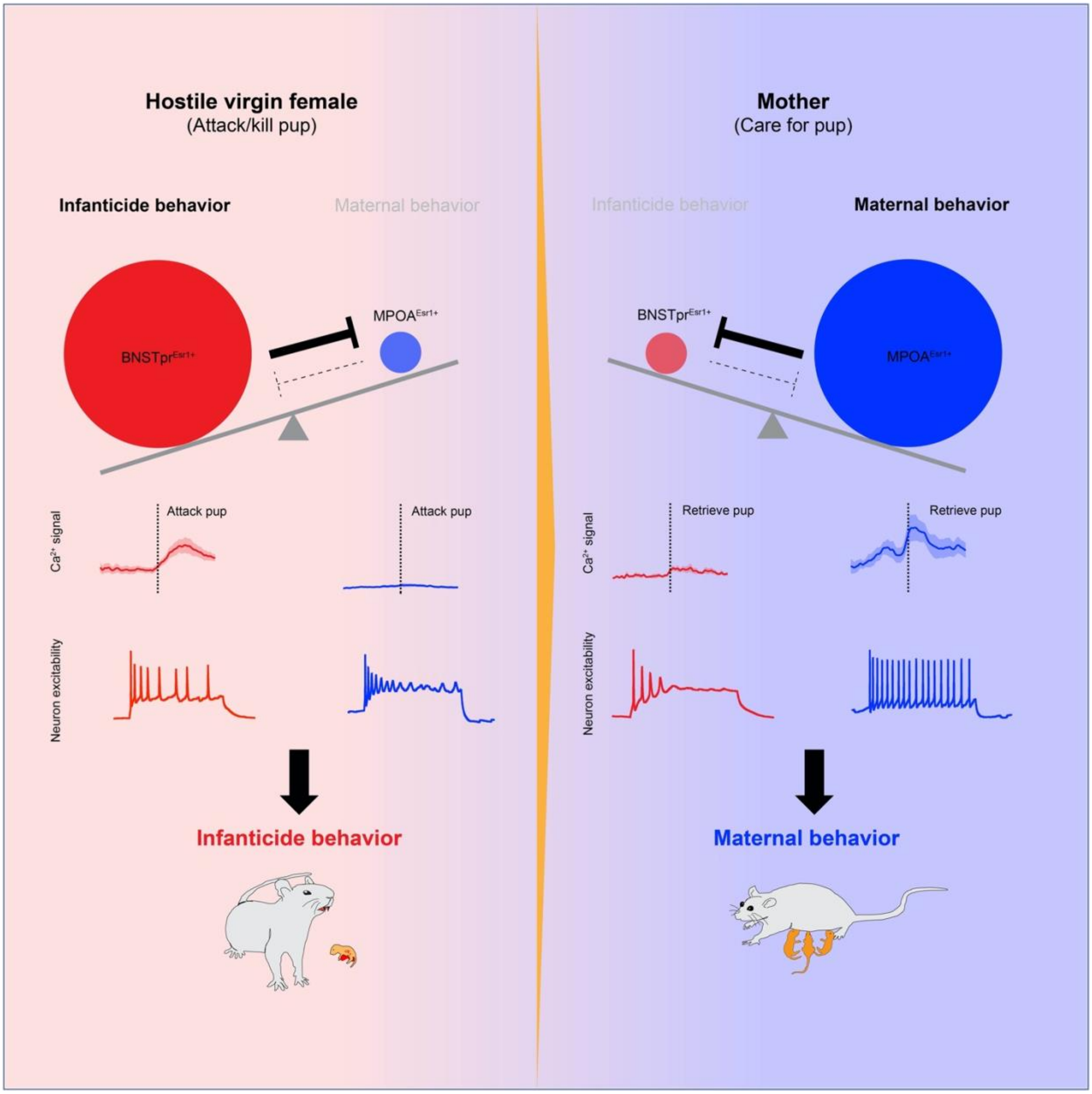


**Extended Data Fig. 8: Schematic showing the antagonistic neural circuits that drive opposing behaviors towards the young in females.**
